# *In silico* study of selected alkaloids as dual inhibitors of β and γ-Secretases for Alzheimer’s disease

**DOI:** 10.1101/2024.09.10.612359

**Authors:** Isreal Ayobami Onifade, Haruna Isiyaku Umar, Abdullahi Tunde Aborode, Aeshah A. Awaji, Ifeoluwapo Deborah Jegede, Bunmi Helen Adeleye, Dorcas Oladayo Fatoba

## Abstract

Alzheimer’s disease (AD) has become ubiquitous as the number of elderly individuals increases and has been conceived as a socioeconomic problem lately. To date, no success is recorded for disease-modifying therapies for AD but only drugs for symptomatic relief exist. Research has been centered on the role of β-amyloid on the pathogenesis of AD, which has led to the development of drugs that target Aβ (β and γ-Secretase inhibitors) to reduce the amount of Aβ formed. However, the existing β and γ-Secretase inhibitors were associated with harmful side effects, low efficacy, and inability to cross the blood-brain barrier. Thus, in this current study, 54 alkaloids from the PhytoHub server (phytohub.eu), and two approved drugs were docked against β-Secretases. Additionally, galantamine and 5 alkaloids with the utmost binding potential with β-secretase were subjected to pharmacokinetics evaluation and docked against γ-secretase. . From our result, 5 compounds displayed for both docking periods, with demissidine, solasodine, tomatidine, and solanidine having better BE than the control drugs. Based on the Pharmacokinetics evaluation, 4 compounds possessed good pharmacokinetic evaluation and biological activities than galantamine. This study suggests that demissidine, solasodine, tomatidine, and solanidine are promising dual inhibitors against β and γ-Secretase proteins *in silico*. However, there is an urgent need to carry out in vitro and in vivo experiments on these new leads to validate the findings of this study.

## 1.0 Introduction

AD is a neurodegenerative disorder characterized by the presence of senile plaques in the brain, which are primarily composed of aggregated amyloid-beta (Aβ) that results from the amyloid precursor protein (APP) (Wilson et al., 2012). Amyloid β (Aβ extracellularly) is produced from the proteolytic degradation of amyloid precursor protein and is one of these proteins (APP). APP is a type 1 transmembrane protein that can be found in a variety of tissues but is most commonly found in neurons. It has been proposed that APP is involved in creating synapses and the healing of neurons (Du et al., 2018). Research focuses on three proteases -α-, β- and γ-Secretases-due to their roles in the modulation and production of Aβ and their potential as drug targets for managing AD (Wilson et al., 2012). This study emphasizes β- and γ-Secretases, as they are involved in producing insoluble Aβ, which forms Aβ plaques.

The common symptoms of Alzheimer’s disease include cognitive dysfunction (such as memory loss and communication difficulties), developmental issues (like depression and hallucinations, often known as the non-markers), and challenges with daily activities (Mendez, 2012; Wilson et al., 2012; Burns et al., 1990). The primary causes of AD are genetic predisposition, age, and abnormal protein accumulation within and outside of neurons (Mendez, 2012). Amyloid β (Aβ) is a protein, produced from the proteolytic breakdown of of APP, a type 1 transmembrane protein predominantly found in neurons and a variety of tissues, and is thought to be involved in synapse formation and neuronal repairs (Du et al., 2018).

The key pathophysiological mechanism of Alzheimer’s disease is the external development of amyloid proteins, known as amyloid plaques. It is also the primary drug benchmark for the inhibitory activity of Aβ yield in AD (Humpel, 2011). It is produced by the protease activity of the β-secretase enzyme. Before the symptoms occur, unexpected changes and improvements relative to the brain’s biochemistry are thought to occur (Lesné et al., 2013). In recent times, more information has accumulated addressing the influence of AD caused by significant neuronal damage and neurodegenerative abnormalities in numerous areas of the brain (Lesné et al., 2013). Amyloid plaque is one of the neuropathological hallmarks of AD. It is mainly comprised of Aβ peptide, a 38 to 43 amino acid peptide derived from the cleavage of the β-amyloid precursor protein (βAPP). This latter protein undergoes two distinct and competing mechanisms, usually referred to as the amyloidogenic and non-amyloidogenic pathways, in which the proteolytic mechanisms are largely dependent on three kinds of secretase enzymes (α-, β-, and γ-secretase) (Eisele et al., 2015). In addition, almost all drugs developed over the years to treat AD were reported to have failed to exhibit the expected efficacy. Furthermore, those targeting these proteases as either enhancers, modulators, or inhibitors were discontinued before phase III trials. Hence, there is an urgent need to discover small molecules from natural sources such as plants to avert or slow down the progression of AD.

Alkaloids are nitrogen-containing plant molecules found majorly among plant families such as *Papaveraceae* (poppies family), *Amaryllidaceae* (amaryllis), *Solanaceae* (nightshades), and *Ranunculaceae* (buttercups). Plant alkaloids exhibit a wide range of biological and pharmacological potentials, including antidiabetic potential (Rasuoli et al., 2020). Due to its antidiabetic potential, we explored some plant alkaloids in this current *in silico* study. Interestingly, two drugs approved by the FDA as cholinesterase inhibitors are from this class of plant molecules (galantamine and rivastigmine) to treat AD (De Strooper, 2010). This suggests the importance of this class of compounds. However, little research has looked at alkaloids as potential inhibitors against β- and γ-Secretases to treat AD. This study therefore used in silico approach to predict the inhibitory properties of alkaloids as potential drug targets against AD.

## 2.0 Materials and Methods

### 2.1 Selection and Preparation of Protein Targets

The 3D structures of β- and γ-Secretases from the Protein Data Bank were retrieved with PDB codes 4LXM (Rueeger et al., 2013) and 7D8X (G. Yang et al., 2021) respectively. Then, they were subjected to cleaning and optimization through the dock prep module of UCSF-Chimera software. Here, the proteins were freed from all heteroatoms such as co-crystallized Ligands and water. Followed by hydrogenation and applying charges using gasteiger charge. Finally, the proteins were minimized under the influence of amber force field 94 fs (Amberff94fs) (Pettersen et al., 2004).

### 2.2 Selection and Optimization of Ligands

86 compounds of alkaloid extraction (Table 1) were selected as our Ligands under investigation with two approved drugs to treat AD from Drugbank (galantamine and rivastigmine). Their 3D conformers were obtained through the phytohub webserver (phytohub.eu) in structure data format (SDF). Open Babel housed within the Python prescription (PyRx) suit was deployed to filter the compounds through their molecular weights (<500), then those having a molecular weight less than 500 were optimized under the influence of the Universal Force field (UFF) and converted to their best energetic and stable configurations.

**Table 1:** Selected alkaloids in food.

| S/No. | Compounds | Phytohub ID | Pubchem CID |
| --- | --- | --- | --- |
|  | 15-decarboxy-betanin | PHUB000416 | 12300103 |
|  | 17-decarboxy-betanin | PHUB000413 | 101333461 |
|  | 17-decarboxy-neobetanin | PHUB000415 | 102401025 |
|  | 2-decarboxy-betanin | PHUB000410 | 131751981 |
|  | 3-Methoxy-tyramine-betaxanthin | PHUB000431 | 135871118 |
|  | 5-Hydroxynorvaline-betaxanthin | PHUB000456 | 246598 |
|  | Alanine-betaxanthin | PHUB001470 | 24779335 |
|  | Arginine-betaxanthin | PHUB000451 | 447276923 |
|  | Asparagine-betaxanthin | PHUB000434 | 447276924 |
|  | Betalamic acid | PHUB000404 | 5281176 |
|  | Betanidin | PHUB000407 | 135449343 |
|  | Betanin | PHUB000405 | 12300103 |
|  | glutamine-betaxanthin | PHUB000432 | 135438599 |
|  | Glycine-betaxanthin | PHUB000444 | 135438598 |
|  | Gomphrenin I | PHUB000429 | 6096868 |
|  | Gomphrenin II | PHUB001477 | 131752748 |
|  | Gomphrenin III | PHUB001476 | 101105498 |
|  | Histamine-betaxanthin | PHUB000446 | 774 |
|  | Histidine-betaxanthin | PHUB000442 | 6274 |
|  | Hylocerenin | PHUB000423 | 101168510 |
|  | Isobetanidin | PHUB000408 | 135612764 |
|  | Isobetanin | PHUB000406 | 6325438 |
|  | Isogomphrenin I | PHUB000430 | 90658633 |
|  | Isogomphrenin II | PHUB001478 | 101105497 |
|  | Isogomphrenin III | PHUB001479 | 101105499 |
|  | Isolampranthin II | PHUB000428 | 157010344 |
|  | Isoleucine-betaxanthin | PHUB000436 | 157010345 |
|  | Isophyllocactin | PHUB000426 | 101168512 |
|  | Lampranthin II | PHUB000427 | 11953909 |
|  | Leucine-betaxanthin | PHUB000435 | 447276973 |
|  | Neobetanin | PHUB000409 | 102401026 |
|  | Phenylalanine-betaxanthin | PHUB000448 | 52923786 |
|  | Phyllocactin | PHUB000425 | 101056997 |
|  | Prebetanin | PHUB000422 | 6325833 |
|  | Proline-betaxanthin | PHUB000447 | 57513848 |
|  | Tryptophan-betaxanthin | PHUB000454 | 136728070 |
|  | Tyramine-betaxanthin | PHUB000440 | 157010370 |
|  | tyrosine-betaxanthin | PHUB000443 | 135887852 |
|  | Valine-betaxanthin | PHUB000450 | 447277025 |
| | γ-Aminobutyric acid-betaxanthin | PHUB000445 | 447277027 |
|  | 13-Hydroxylupanine | PHUB001464 | 250873 |
|  | Adenine | PHUB001480 | 190 |
|  | Albine | PHUB000825 | 442936 |
|  | Calystegin A3 | PHUB000826 | 442999 |
|  | Calystegin B2 | PHUB000827 | 443000 |
|  | Gentianine | PHUB000828 | 354616 |
|  | Harman | PHUB000829 | 5281404 |
|  | Harmine | PHUB000830 | 5280953 |
|  | Hyoscyamine | PHUB000831 | 154417 |
|  | lupanine | PHUB001465 | 91471 |
|  | lupinine | PHUB001467 | 91461 |
|  | Multiflorine | PHUB001468 | 6918763 |
|  | Perlolyrine | PHUB000832 | 160179 |
|  | Physoperuvine | PHUB000833 | 443008 |
|  | Salsolinol ((-)-) | PHUB000834 | 91588 |
|  | Scopolamine | PHUB000835 | 3000322 |
|  | Sparteine | PHUB001469 | 644020 |
|  | Trigonelline | PHUB001378 | 5570 |
|  | 1,3-dimethyluric acid | PHUB002107 | 70346 |
|  | Caffeine | PHUB000789 | 2519 |
|  | Convicine | PHUB000793 | 88000 |
|  | Dihydrozeatin | PHUB000795 | 32021 |
|  | Theobromine | PHUB000790 | 5429 |
|  | Theophylline | PHUB000791 | 2153 |
|  | Vicine | PHUB000792 | 135413566 |
|  | Zeatin (trans-) | PHUB000794 | 449093 |
|  | Arecaidine | PHUB000836 | 10355 |
|  | Arecoline | PHUB000837 | 2230 |
|  | Carpaine | PHUB000838 | 442630 |
|  | Cucurbitine | PHUB000839 | 442634 |
|  | Fagomine | PHUB000840 | 72259 |
|  | Guvacine | PHUB000841 | 3532 |
|  | Pelletierine ((-)-) | PHUB000842 | 92987 |
|  | Piperine | PHUB000843 | 638024 |
|  | Trichostachine | PHUB000845 | 636537 |
|  | Chaconine (alpha-) | PHUB000846 | 442971 |
|  | Dehydrotomatine | PHUB000847 | 131751243 |
|  | Demissidine | PHUB000848 | 101379 |
|  | Demissine | PHUB000849 | 442975 |
|  | Solamargine | PHUB000850 | 73611 |
|  | Solanidine | PHUB000851 | 65727 |
|  | Solanine (alpha-) | PHUB000852 | 159233 |
|  | Solasodine | PHUB000853 | 442985 |
|  | Solasonine | PHUB000854 | 119247 |
|  | Tomatidine | PHUB000855 | 65576 |
|  | Tomatine | PHUB000856 | 28523 |

### 2.3 Virtual Docking simulation of β-secretase and ligands

Before the docking simulation, the protocol for docking was validated by re-docking the co-crystallized ligand into its binding region in β-secretase to regenerate a similar binding post like that in the X-ray crystallized structure of β-secretase (4LXM). Auto dock vina was used for the docking and PyMOL 2.4 was used to align the two complex structures, consequently obtaining the root mean square deviation (RMSD). The virtual docking of all selected ligands against β-secretase was achieved by employing the flexible procedure of docking previously used (Umar et al., 2021). PyRx 0.8, a suite integrated with Auto Dock Vina, was applied for the molecular docking simulation (Trott & Olson, 2010). The specific target site for β-secretase corresponding to the binding region used in APP processing was adjusted using the grid box with dimensions (23.1362 x 19.9375 x 23.7691) Å, and the centre was attuned based on the site of binding in the protein consisting of the following amino acids; Asp228, Asp32, Gln73, Thr72 and Gln12 (Rueeger et al., 2013). The compounds with better Binding energy than the control drugs at the end of the experiment were subjected to drug-likeness screening using the SwissAdme server (Daina et al., 2017) and molecular interaction analysis with the aid of PyMOL© Molecular Graphics (version 2.4, 2016, Shrodinger LLC) and LigPlot^+^. The generated compounds are regarded as potential hits.

### 2.4 Molecular Docking simulation of γ-secretase and hit Ligands

Similar to the above steps, the molecular docking was carried out between γ-secretase and hit compounds with good pharmacokinetic attributes based on the outcome of drug-likeness screening via the SwissAdme server (Daina et al., 2017). However, comparing the molecular interactions in the co-crystallized and re-docked complexes were carried out for the protocol validation. Furthermore, the specific target site for γ-secretase corresponding to the binding region exercised in APP processing was attuned via the grid box with dimensions (26.514 x 24.0892 x 25.4268) Å, and the centre was acclimated based on the catalytic site binding in the target comprising of the listed amino acids; Leu381, Pro433, Ala434, Leu435, Lys380, Asp257, Leu432, Tyr288, Asp385, Ser289, Val261, Val272, Ser290, Gly382, Gly384, Leu282, Ile143, Ile253, Tyr256, Phe388, Thr421, Leu425 and Val379 (G. Yang et al., 2021).

### 2.5 In silico pharmacokinetics screening

Hit compounds against the secretase enzymes were screened for their pharmacokinetics and physicochemical properties with the assistance of the SwissAdme server (Daina et al., 2017) after uploading their canonical smiles.

### 2.6 Molecular Dynamics Simulation analyses

The hit alkaloids were arranged after the thorough computational virtual screening, molecular docking analysis, and ADMET analysis. The selected protein-ligand complexes rigid binding stability and molecular pattern in the physiological sphere were validated via molecular dynamic (MD) simulation evaluation of 100 ns (Bappy *et al.,* 2023, Danazumi and Umar, 2023, Moin *et al.,* 2022). The Schrodinger’s Desmond module was deployed for MD simulation investigations (Bowers *et al.,* 2006). The analysis begins with introducing the best-docked poses of demissidine, solanidine, solasidine, Tomatidine, and galantamine ligand-protein complexes for the two secretases, we are probing in this study, to Desmond module in the proper file format. The protein preparation wizard (Sastry *et al.,* 2013) was utilized for selected ligand and protein structural optimization by removing and repairing steric hindrances, disordered and undesirable atomic geometries.

The solvated water–soaked MD simulation system was generated using the Desmond System Builder tool. The intermolecular interaction potential 3points transferable (TIP3P) model was turned to account for solvated model inception. An orthorhombic simple point charge water box of 8Å beyond to any boundary of the protein-ligand complex was precisely invented. The OPLS-3e force field was deployed for energy minimization (Harder *et al.,* 2016). The simulated system complex charges were kept at neutral, and isosmotic state by adding desirable amount of the counter ions. A total 59.446 mM, 50.817 mM sodium (Na^+^) and chloride (Cl^-^) ions are added into the simulation system. Then, the simulation system was equilibrated. The NPT ensemble at 300 K temperature and 1.0 bar pressure coupled with known Berendsen coupling protocol is set off for system equilibration. The SHAKE algorithm was deployed for H-bond geometry constrained analysis as a time function. The non-bonded energy computations, particularly long-range electrostatic energy, were computed with the Particle Mesh Ewald (PME) algorithm. The simulation was set for 100 ns and 1000 simulation trajectories frames were retained for the computation of root-mean square fluctuation (RMSF), root-mean square deviation (RMSD), protein-ligand contacts, and interactions fractions (%). The in-built Desmond interaction diagram tool was used for plot and figure generation.

## 3.0 Results and Discussion

### 3.1 Results

The present *in silico* studies aimed at discovering potential dual inhibitors against two key enzymes involved with amyloid protein processing called β and γ-secretase (3D structures provided in Figure 1) from 86 alkaloids sourced from Phytohub server. Out of these 86 compounds (Table 1), 52 were found to have molecular weights less than 500 through the open babel module in PyRx suit 0.8. consequently, these 52 compounds and two controls (2D structures provided in Figure 2) were docked against the active site of β-secretase through autodock vina. We observed that binding energies (BE) that results are from -4.8 to -9.4 kcal/mol where the two control drugs (galantamine and rivastigmine) have BE values of -7.4 and -6.1 kcal/mol, respectively. Five (5) compounds among the 52 alkaloids displayed the utmost BE while 15 alkaloids had BE below that of the two control drugs. The details of these findings are presented in Table 2.

**Figure 1.**
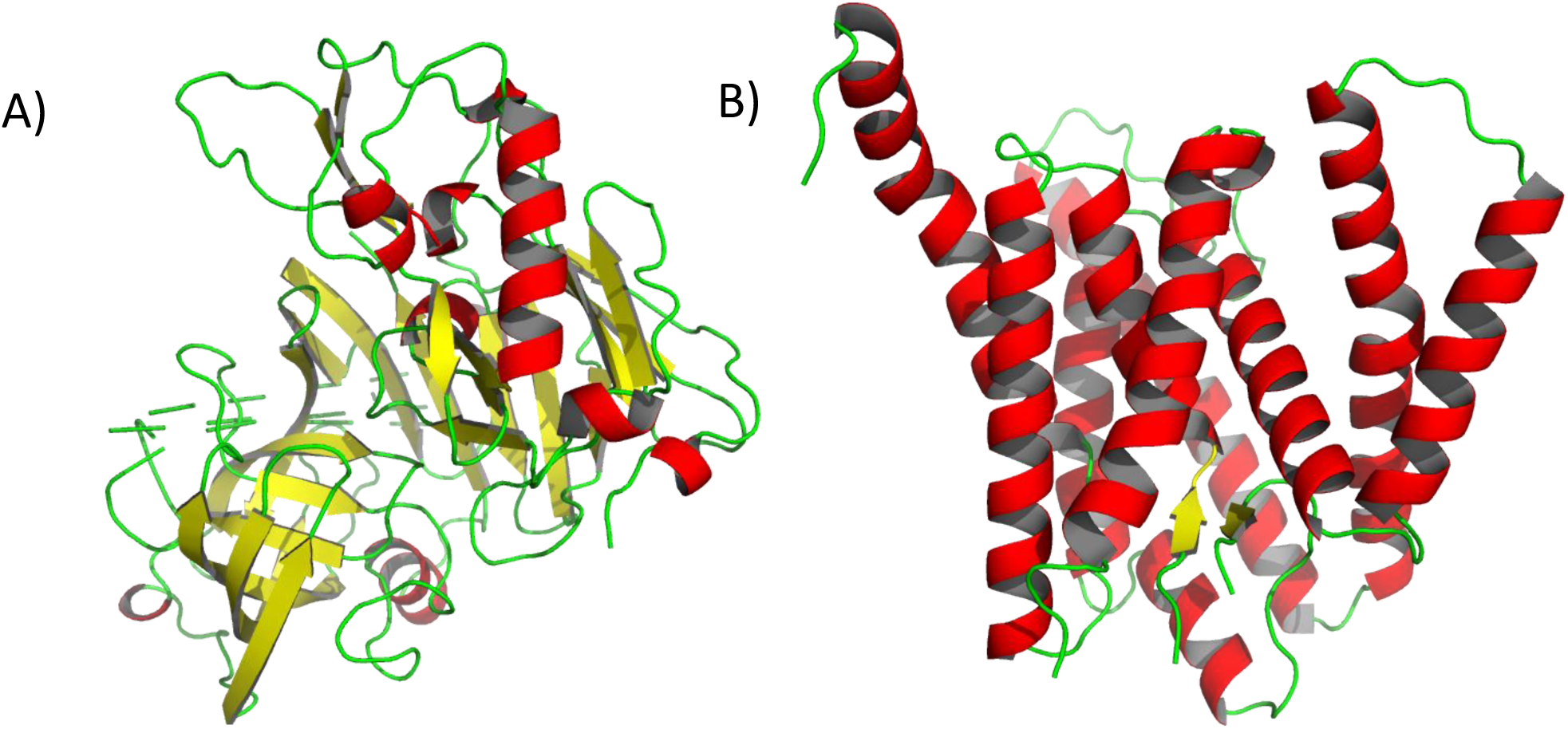
3D structures of Secretases used in our current in silico study retrieved from the protein data bank server. A) β-secretase (4LXM, resolved at 2.30 Å). B) γ-secretase (7D8X, resolved at 2.60 Å)

**Figure 2.**
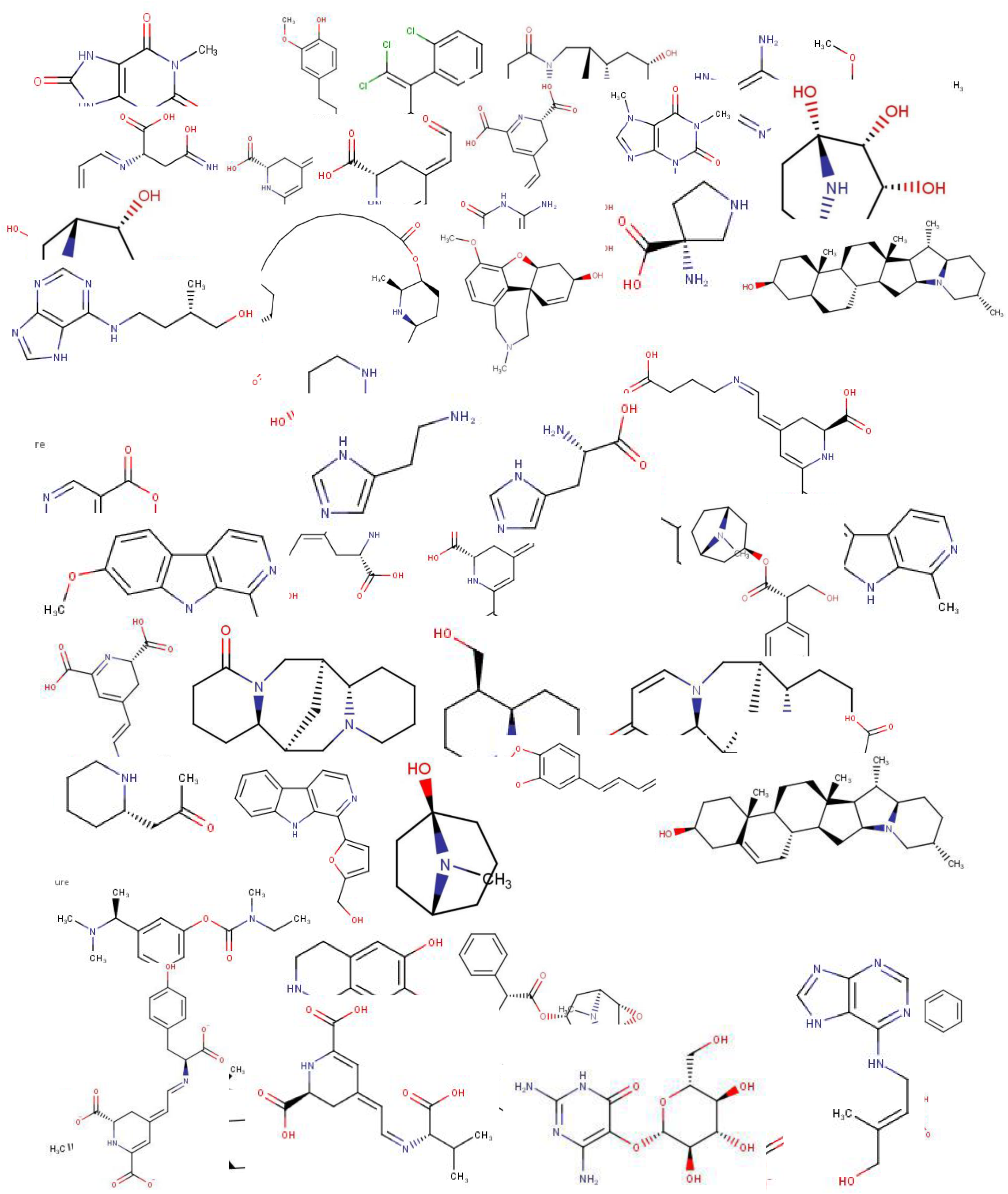
2D structures of alkaloid compounds docked against secretase.

**Table 2.**
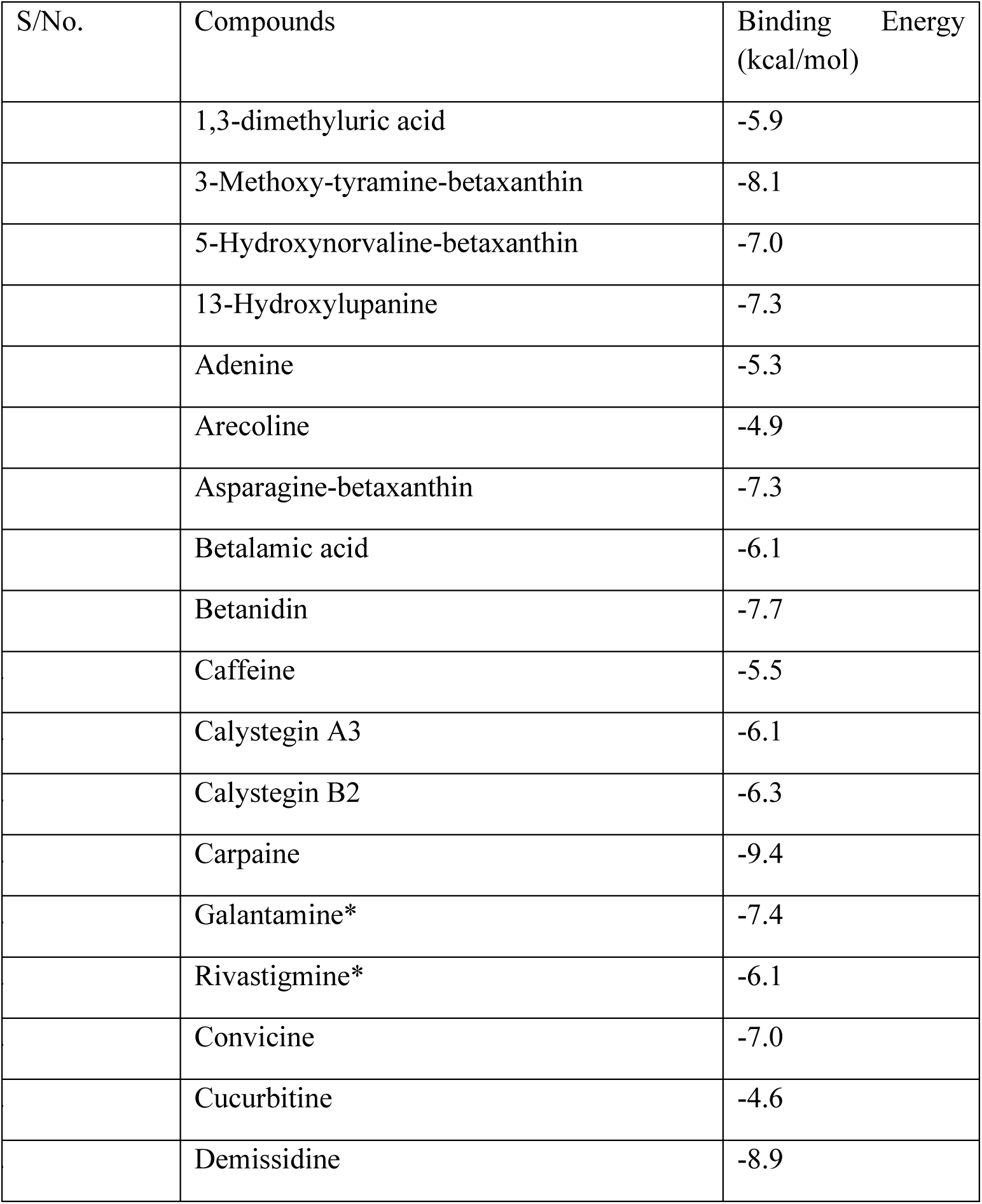
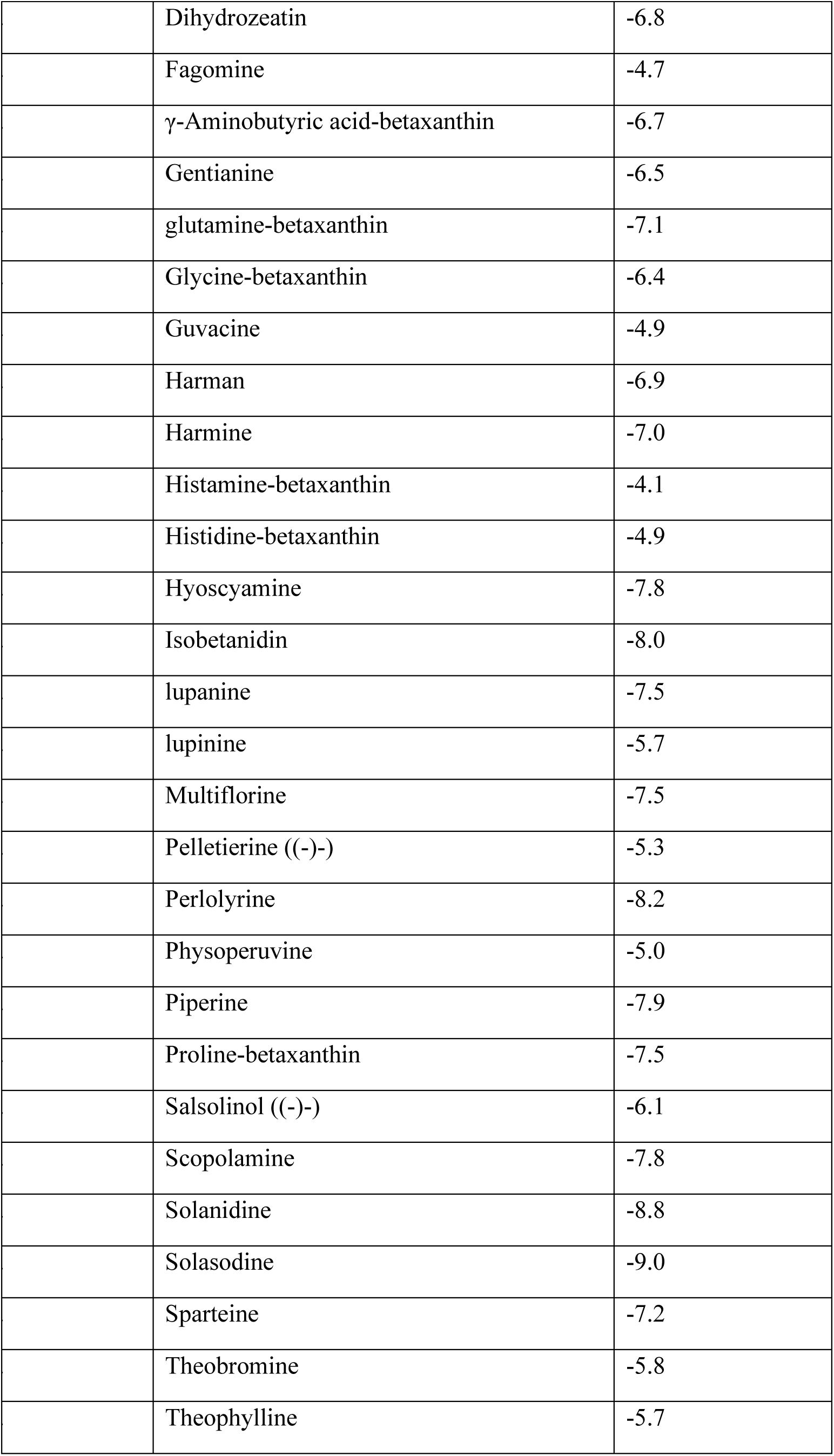
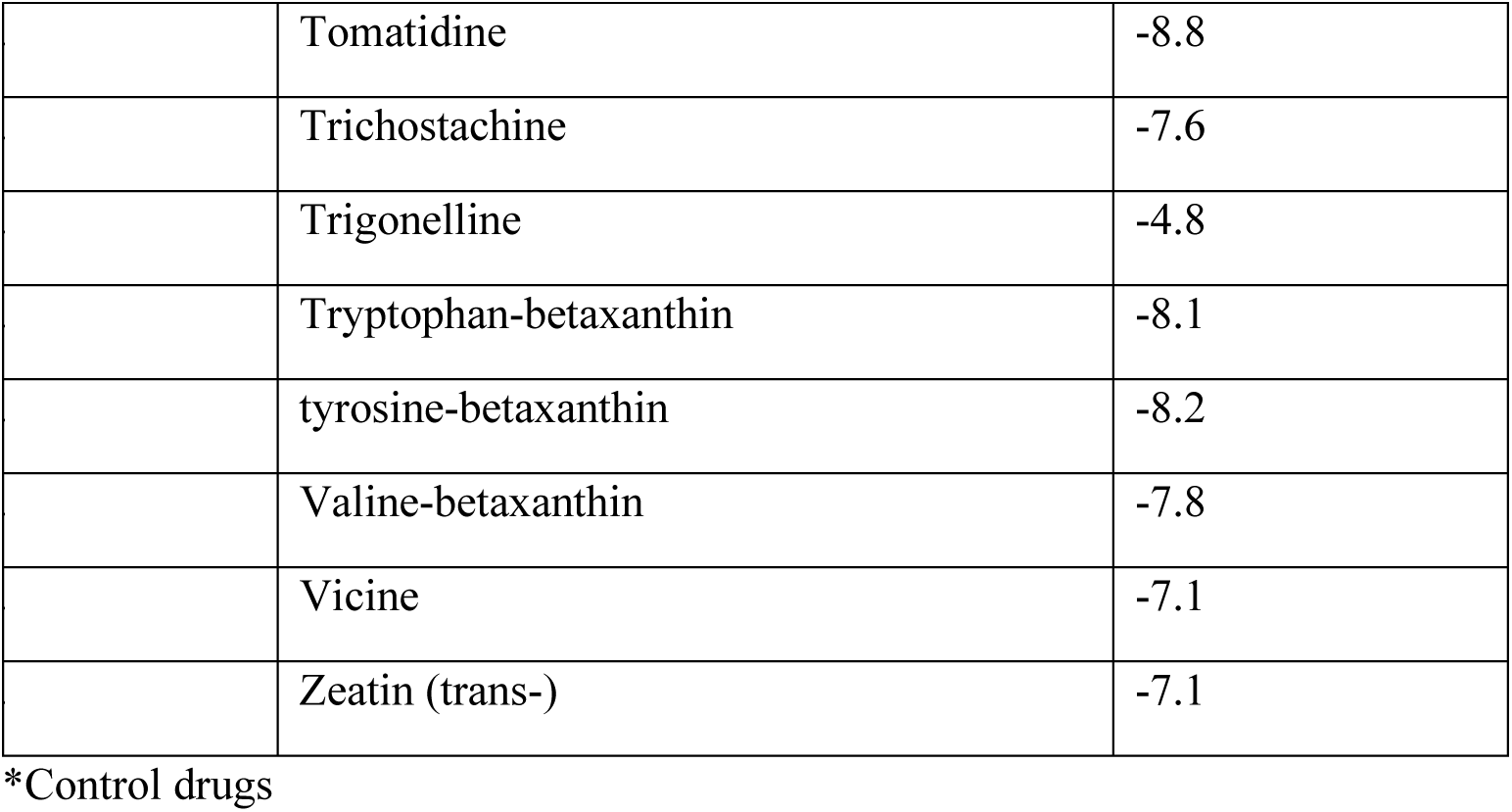
Binding energy (kcal/mol) of compounds docked against β-secretase.

The best control drug and 5 alkaloids with the utmost binding potential with β-secretase were subjected to evaluating drug-likeness and medicinal chemistry properties using the SwissAdme server. The results are presented in Tables 3 and 4. In addition, the molecular interaction analysis was assessed using PyMOL molecular visualization and LigPlot^+^ software for 3D and 2D presentations respectively (Figure 3). The analysis of the protocol validation showed our docking protocol was okay for use (Figure 3a). The binding poses of alkaloids and galantamine were observed to take a similar spot in the binding region of β-secretase (Figure 3b). Figures 3c to 3h present the 2D interaction analysis of compounds with amino acid residues of the target protein.

**Figure 3.**
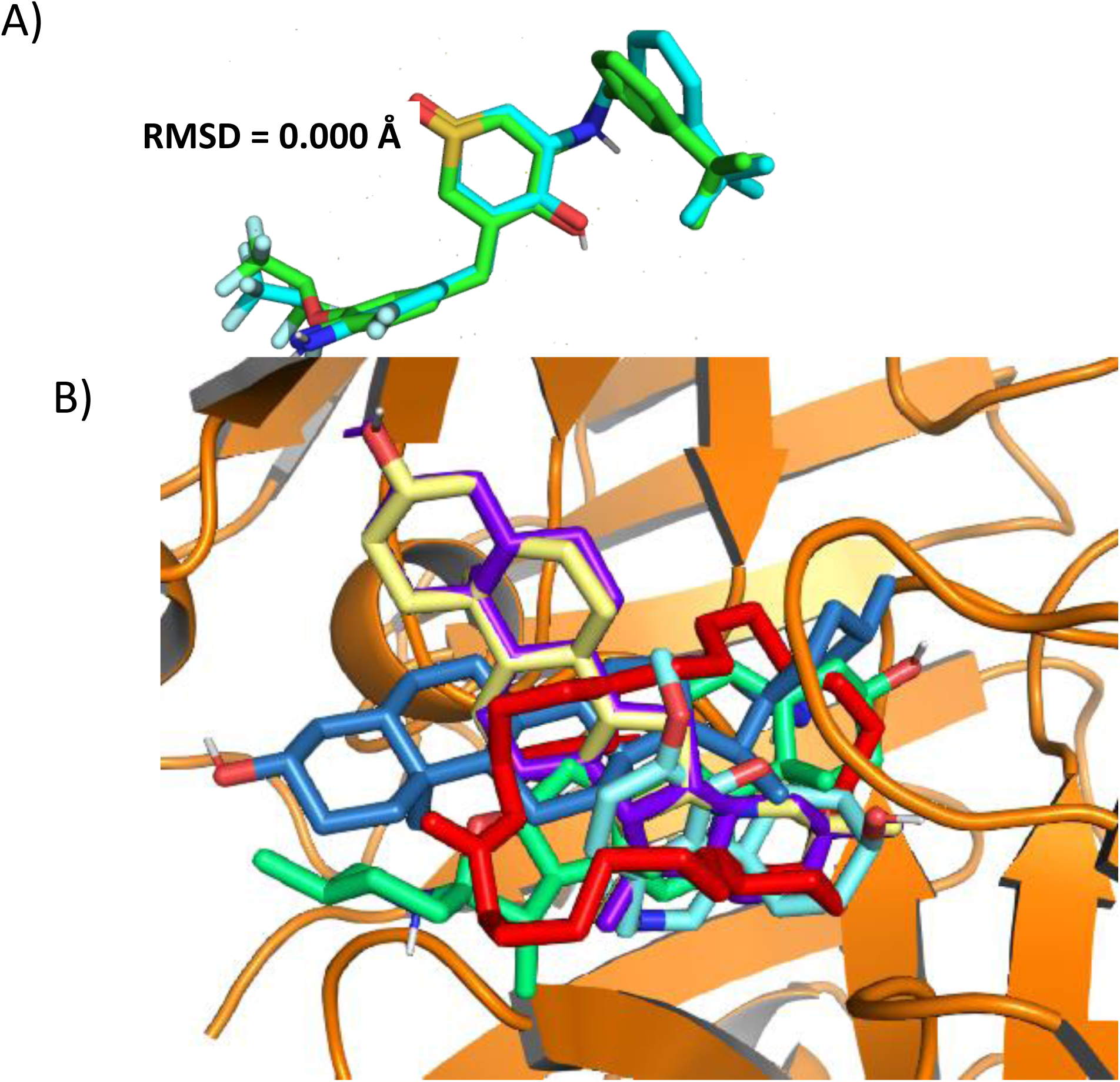
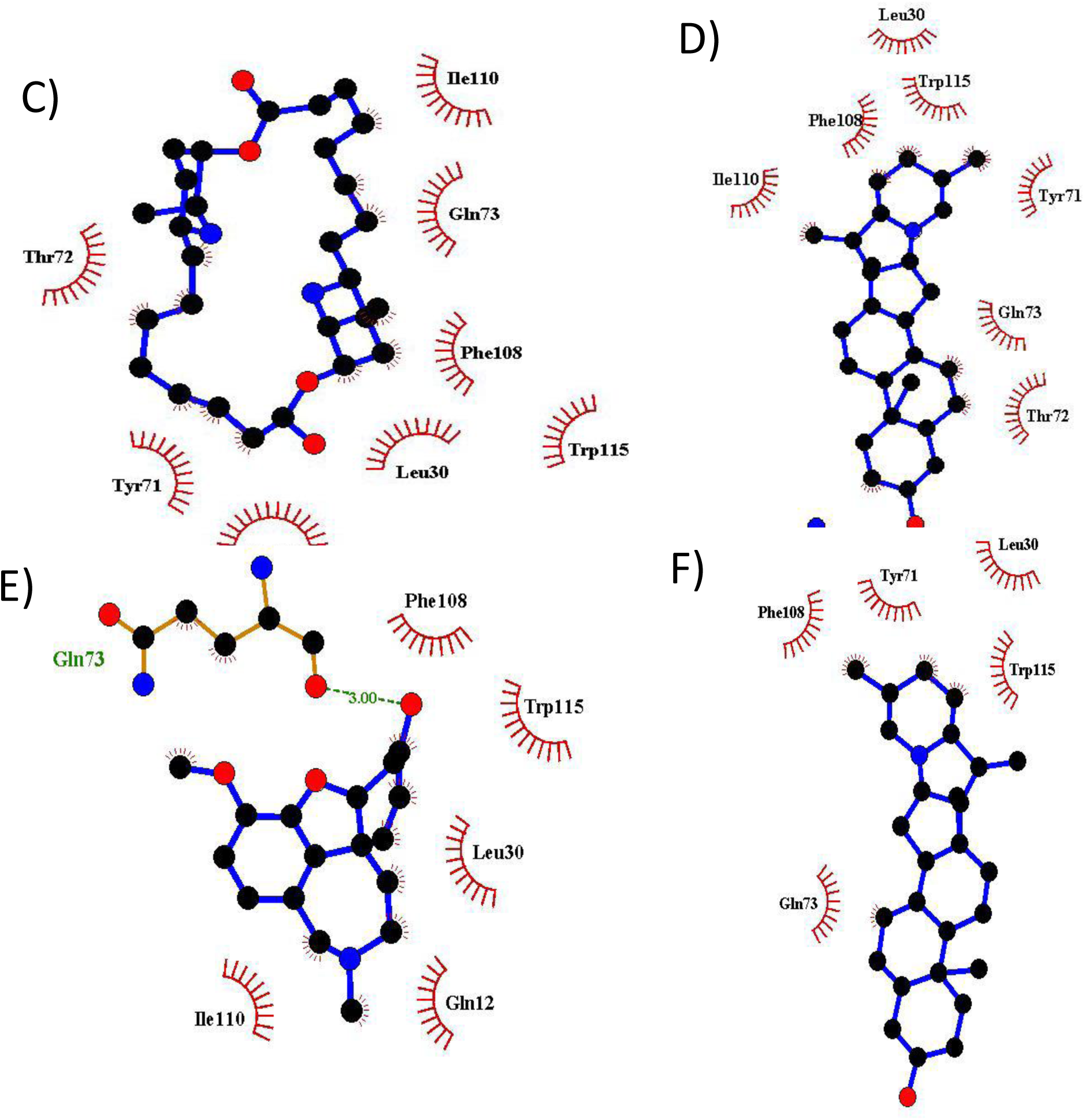
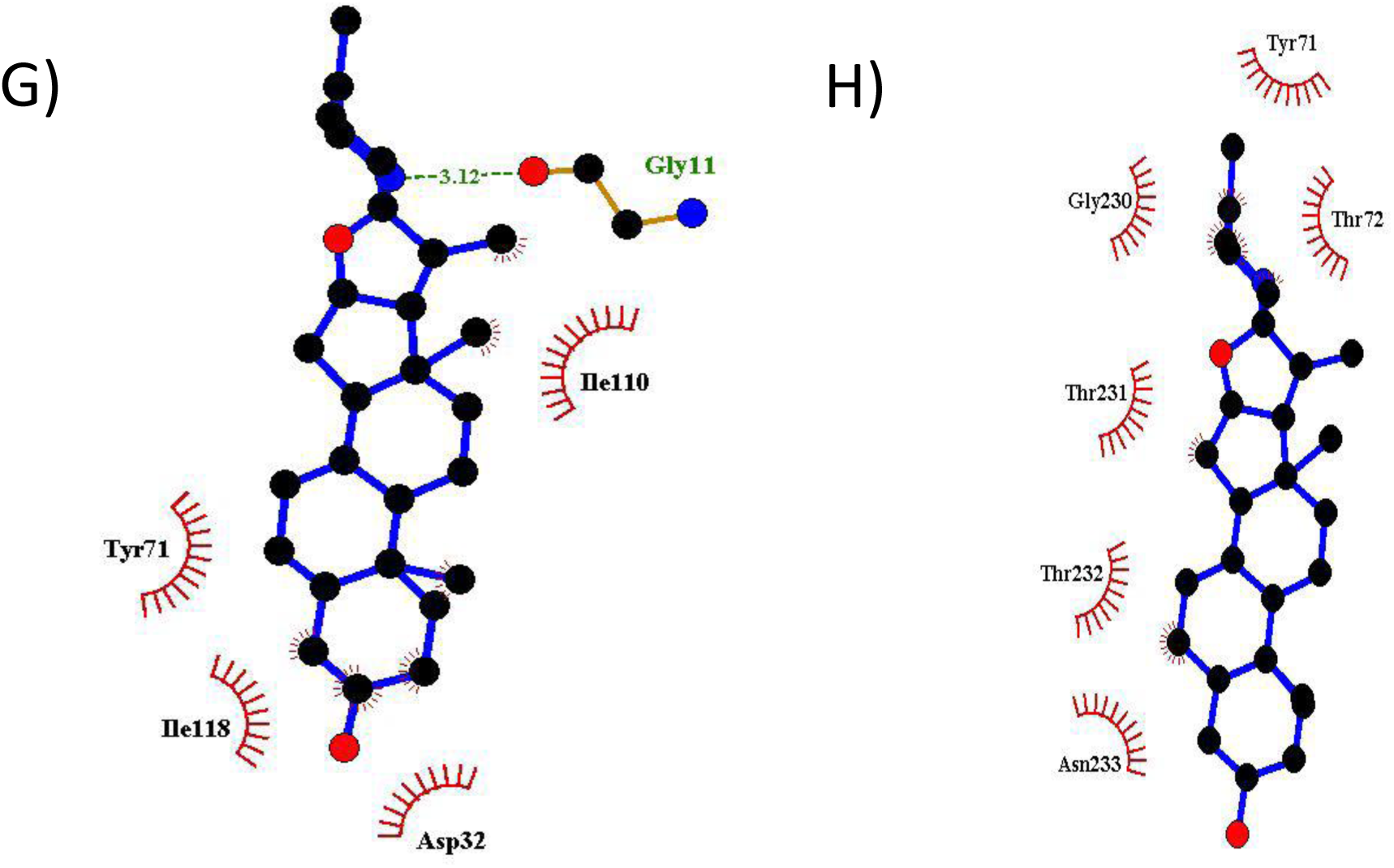
Virtual docking of some alkaloids against β-secretase utilizing Auto Dock Vina, PyMOL and LigPlot^+^ softwares based on the methodology. A) Validation of virtual docking protocol. This process is crucial for the enhancement of the reliability of a computational docking experiment. Comparing the binding postures of co- crystallised ligand (green) with re-docked ligand (cyan) in stick representation. Molecular docking protocol exactly regenerated the binding conformation of crystallographically determined protein-ligand complex with an RMSD value of 0.000 Å. B) The3D binding postures of hit compounds showing their posregiones in the binding site of β- secretase using PyMOL software. Carpaine (red), demissidine (purple), galantamine (cyan), solanidine (yellow), solasidine (green) and tomatidine (blue) were found to occupy similar spot within the binding region of β-secretase. The 2D molecular interaction analysis was carried out through LigPlot^+^ software and snapshots were taken appropriately. (C) Carpaine (D) Demissidine (E) Galantamine (F) Solanidine (G) Solasidine (H) Tomatidine. Interaction analysis shows hydrogen bond (green dashed lines), and hydrophobic interaction (red curved lines) as compounds (purple) interact with the amino acid residues in the catalytic domain of β-secretase.

**Table 3:** Druglikeness evaluation of hit compounds from the virtual docking involving β-secretase and 54 compounds.

| Molecule | MW | #H-bond acceptors | #H-bond donors | Consensus Log P | Fraction Csp3 |
| --- | --- | --- | --- | --- | --- |
| Carpaine | 478.71 | 6 | 2 | 4.64 | 0.93 |
| Demissidine | 399.65 | 2 | 1 | 5.24 | 1 |
| Solanidine | 397.64 | 2 | 1 | 5.02 | 0.93 |
| Solasodine | 413.64 | 3 | 2 | 4.67 | 0.93 |
| Tomatidine | 415.65 | 3 | 2 | 4.9 | 1 |
| Galantamine* | 287.35 | 4 | 1 | 1.91 | 0.53 |

\*Control drug

**Table 4.**
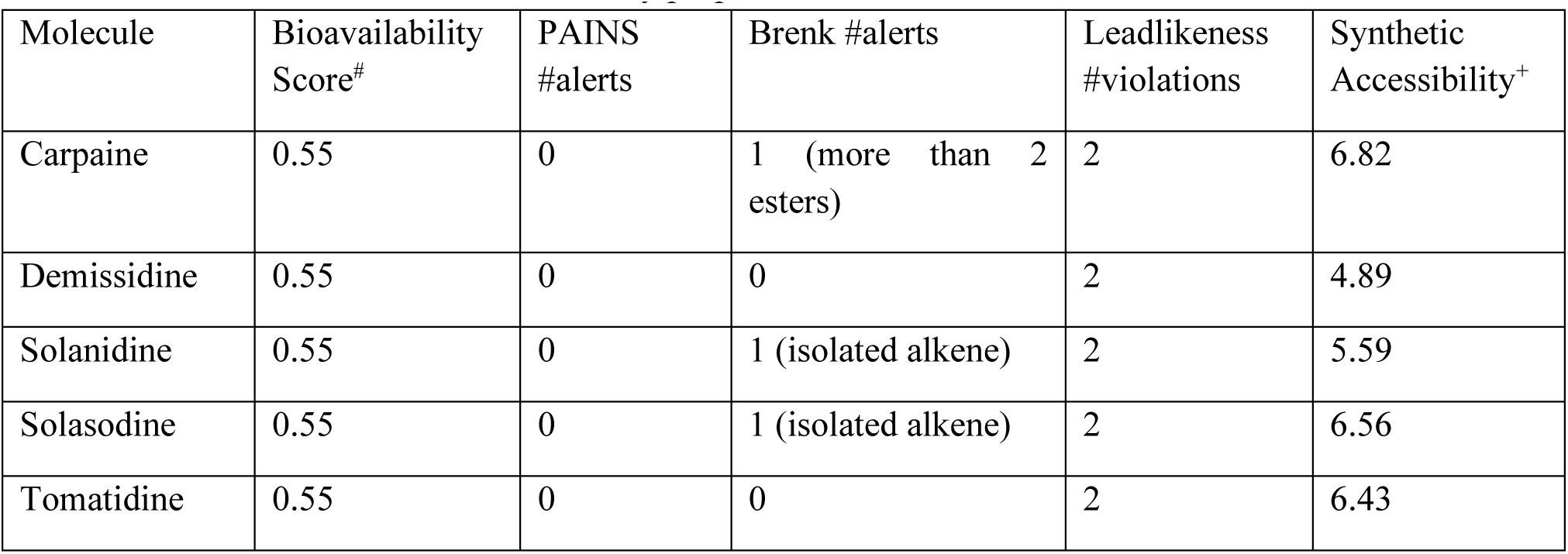
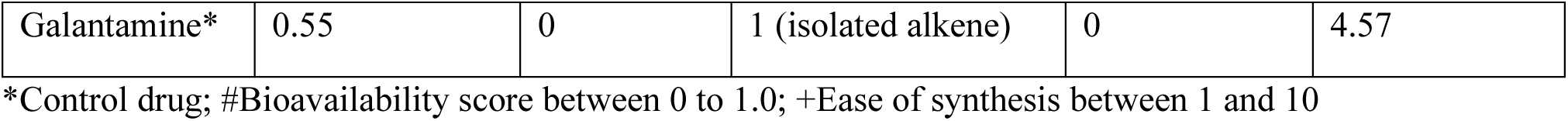
Prediction of the medicinal chemistry properties via the SwissAdme server.

| Molecule | Bioavailability Score<sup>#</sup> | PAINS #alerts | Brenk #alerts | Leadlikeness #violations | Synthetic Accessibility<sup>+</sup> |
| --- | --- | --- | --- | --- | --- |
| Carpaine | 0.55 | 0 | 1 (more than 2 esters) | 2 | 6.82 |
| Demissidine | 0.55 | 0 | 0 | 2 | 4.89 |
| Solanidine | 0.55 | 0 | 1 (isolated alkene) | 2 | 5.59 |
| Solasodine | 0.55 | 0 | 1 (isolated alkene) | 2 | 6.56 |
| Tomatidine | 0.55 | 0 | 0 | 2 | 6.43 |
| Galantamine* | 0.55 | 0 | 1 (isolated alkene) | 0 | 4.57 |

\*Control drug; #Bioavailability score between 0 to 1.0; +Ease of synthesis between 1 and 10

We carried out the second docking against γ-secretase and the five compounds, galantamine and co-crystallized Ligand from the protein data bank. The results obtained here (Table 5) were promising as we observed that demissidine, solasodine, tomatidine and solanidine have BE of -10.2, -10.4, -11.1 and -10.2 kcal/mol respectively, outsmarting the co-crystallized Ligand and galantamine (-9.3 and -7.2 kcal/mol respectively). The molecular interaction analysis and binding poses are provided in Figure 4 similarly done as Figure 3. In a similar light, the *in silico* pharmacokinetic properties were evaluated using the SwissAdme server where properties such as gastrointestinal absorption, blood-brain barrier permeability, substrate of p-glycoprotein, and inhibitors of cytochrome P450 isoforms and excretion through the skin (log Kp). These outcomes are presented in Table 6 which summarily indicates that our compounds are better pharmacokinetically than galantamine. The in-built Desmond interaction diagram tool was used for plot and figure generation.

**Figure 4.**
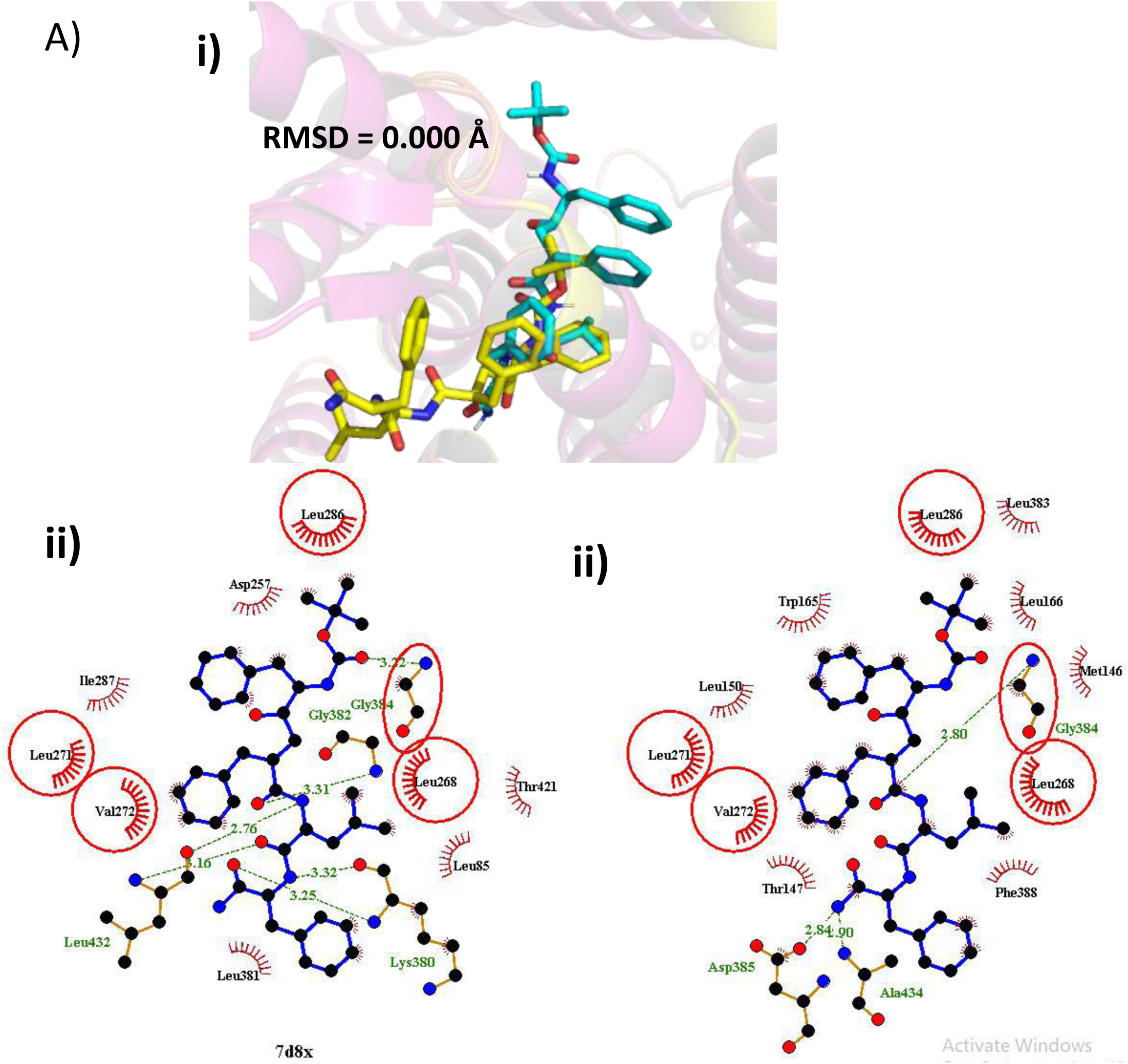
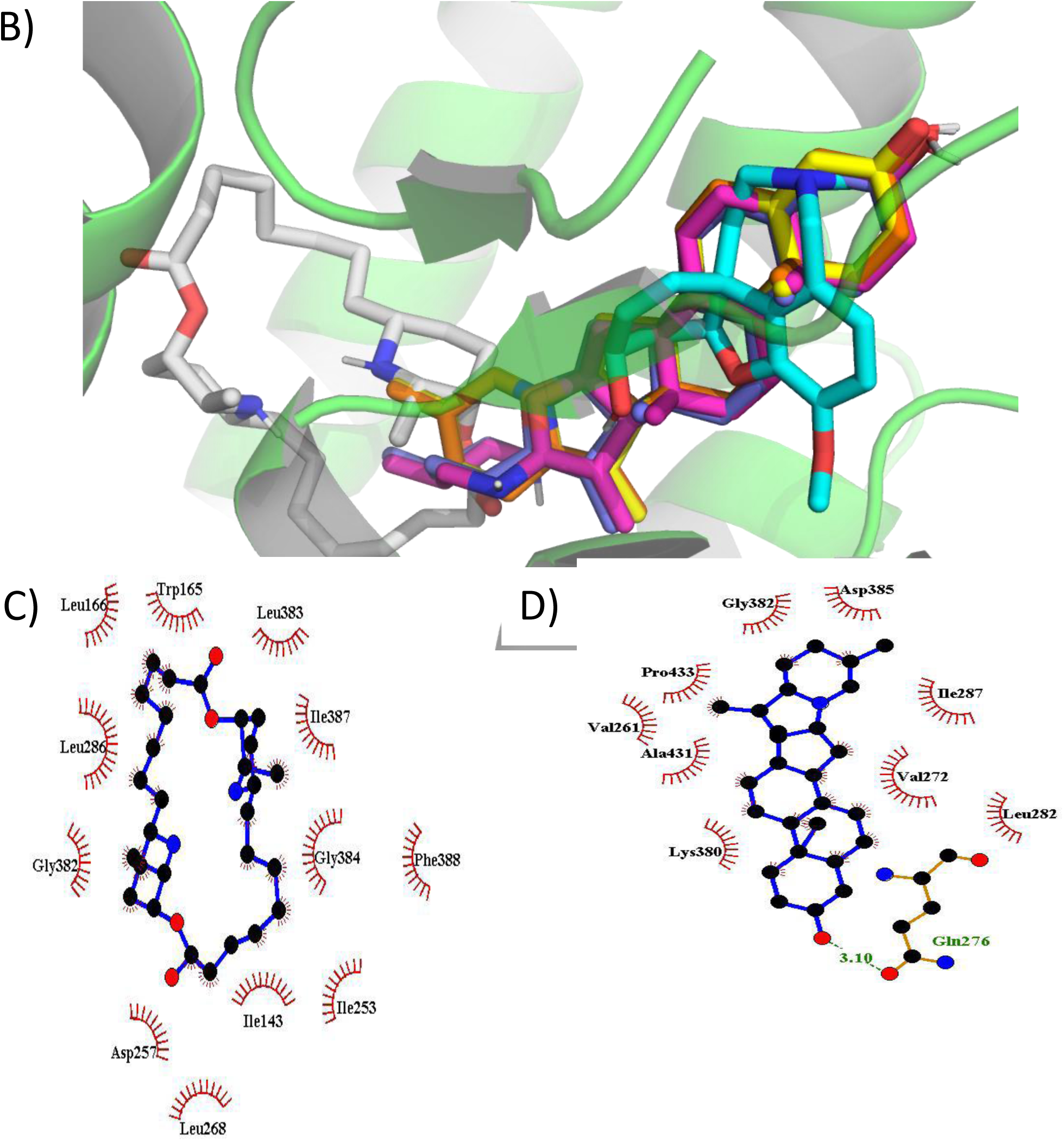
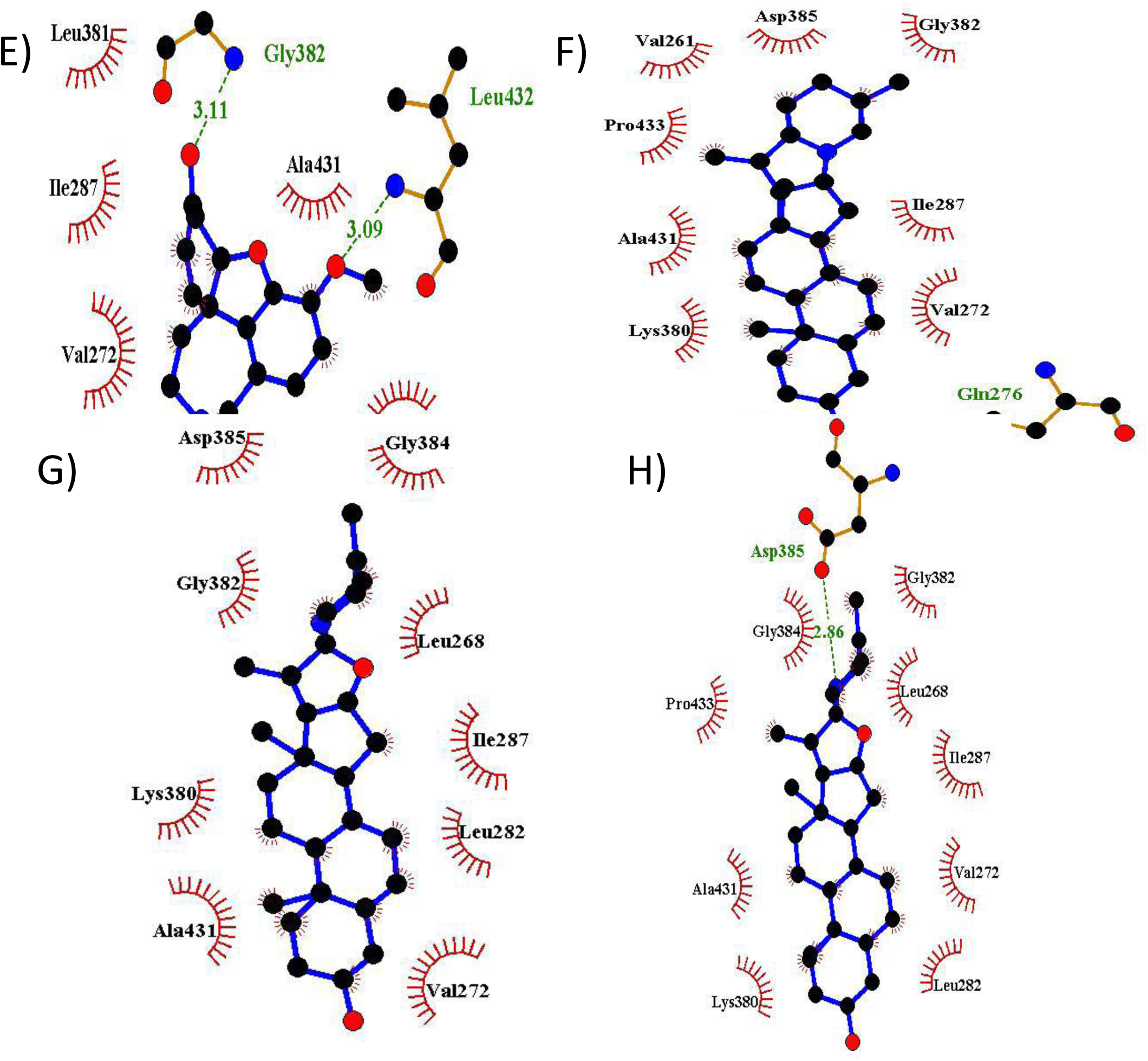
Molecular docking simulation of Hit alkaloids against γ-secretase with the aid of Auto Dock Vina, PyMOL and LigPlot^+^ softwares according to the methodology. A) Validation of docking simulation protocol. i) The binding configurations of co-crystallised ligand (yellow) with re-docked ligand (cyan) in stick illustration. Amino acid residues involved in the atoms of the Ligands in questions are rendered via LigPlot^+^ ii) Co-crystalized L685,458 iii) Re-docked L685,458. B) The 3D binding positions of hit compounds showing their positions in the binding site of γ-secretase using PyMOL software. Carpaine (grey), demissidine (orange), galantamine (cyan), solanidine (yellow), solasidine (blue) and tomatidine (purple) were found to occupy similar spot within the binding region of γ-secretase. The 2D molecular interaction analysis was carried out through LigPlot^+^ software and snapshots were taken appropriately. (C) Carpaine (D) Demissidine (E) Galantamine (F) Solanidine (G) Solasidine (H) Tomatidine. Interaction analysis shows hydrogen bond (green dashed lines), and hydrophobic interaction (red curved lines) as compounds (purple) interact with the amino acid residues in the catalytic domain of γ-secretase.

**Figure 5.**
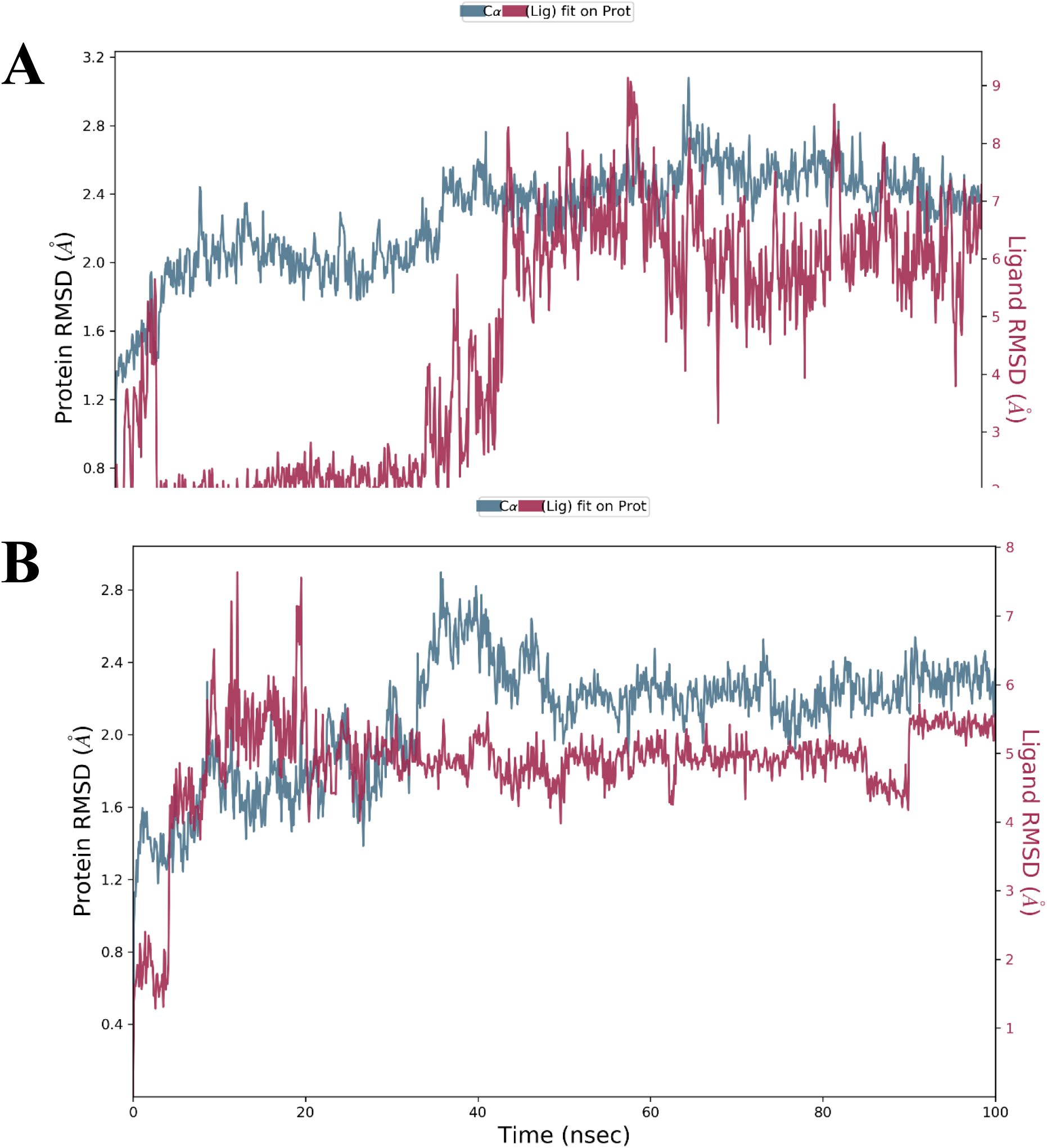
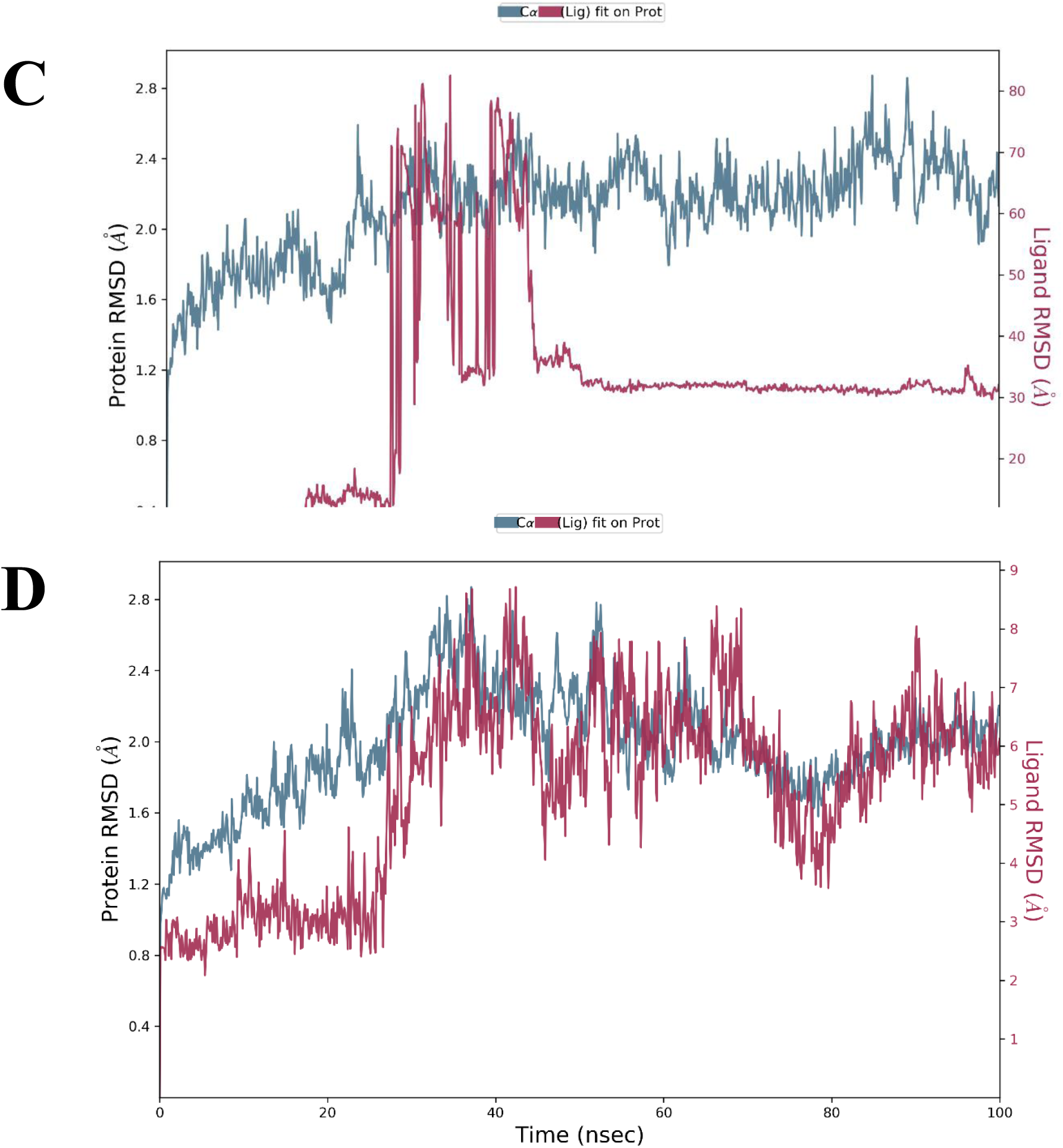
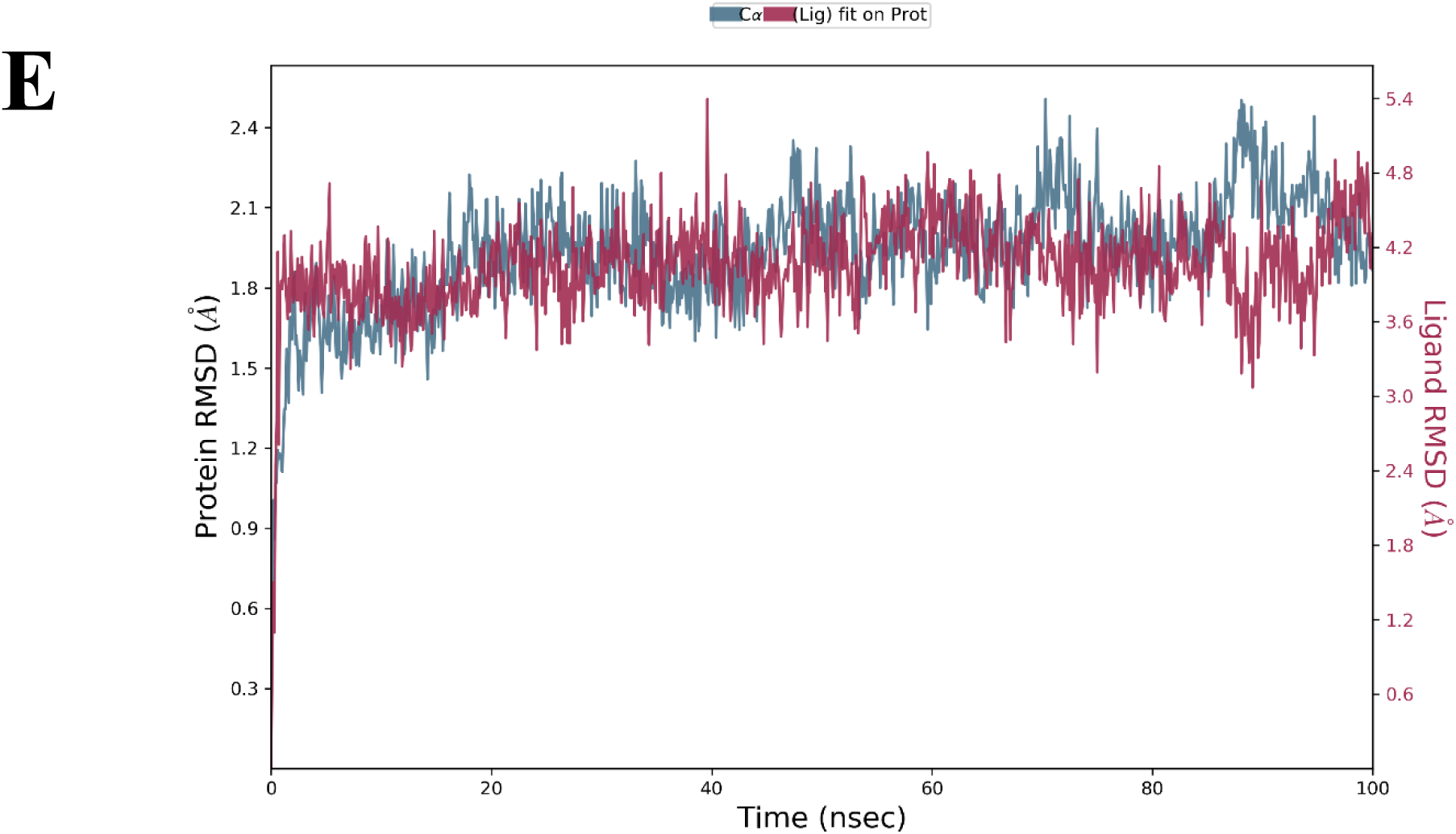
Root Mean Square Deviation of Hit Compound- β-Secretase complexes. A) Demissidine B) Galantamine C) Solanidine D) Solasidine E) Tomatidine.

**Figure 6.**
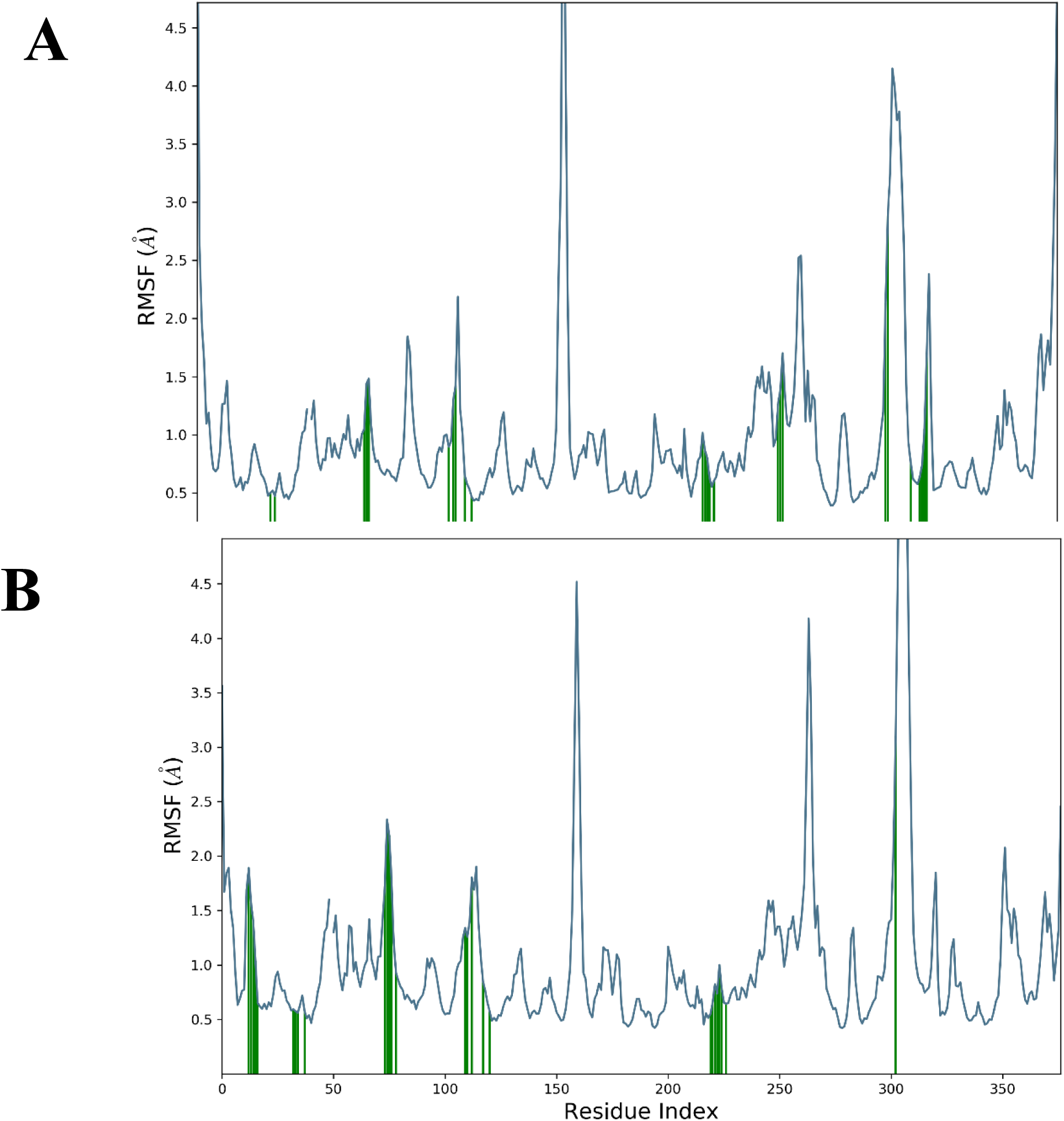
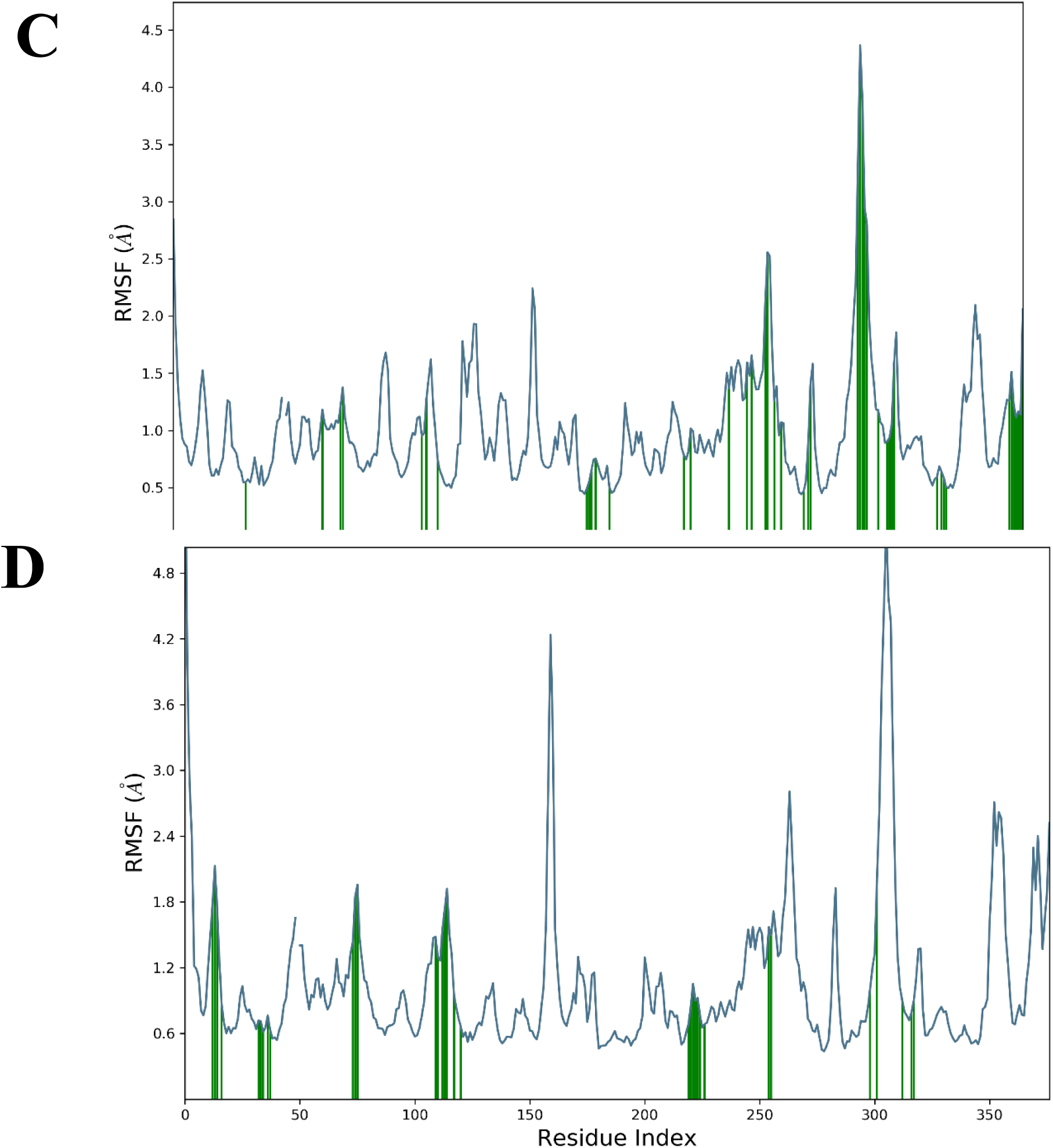
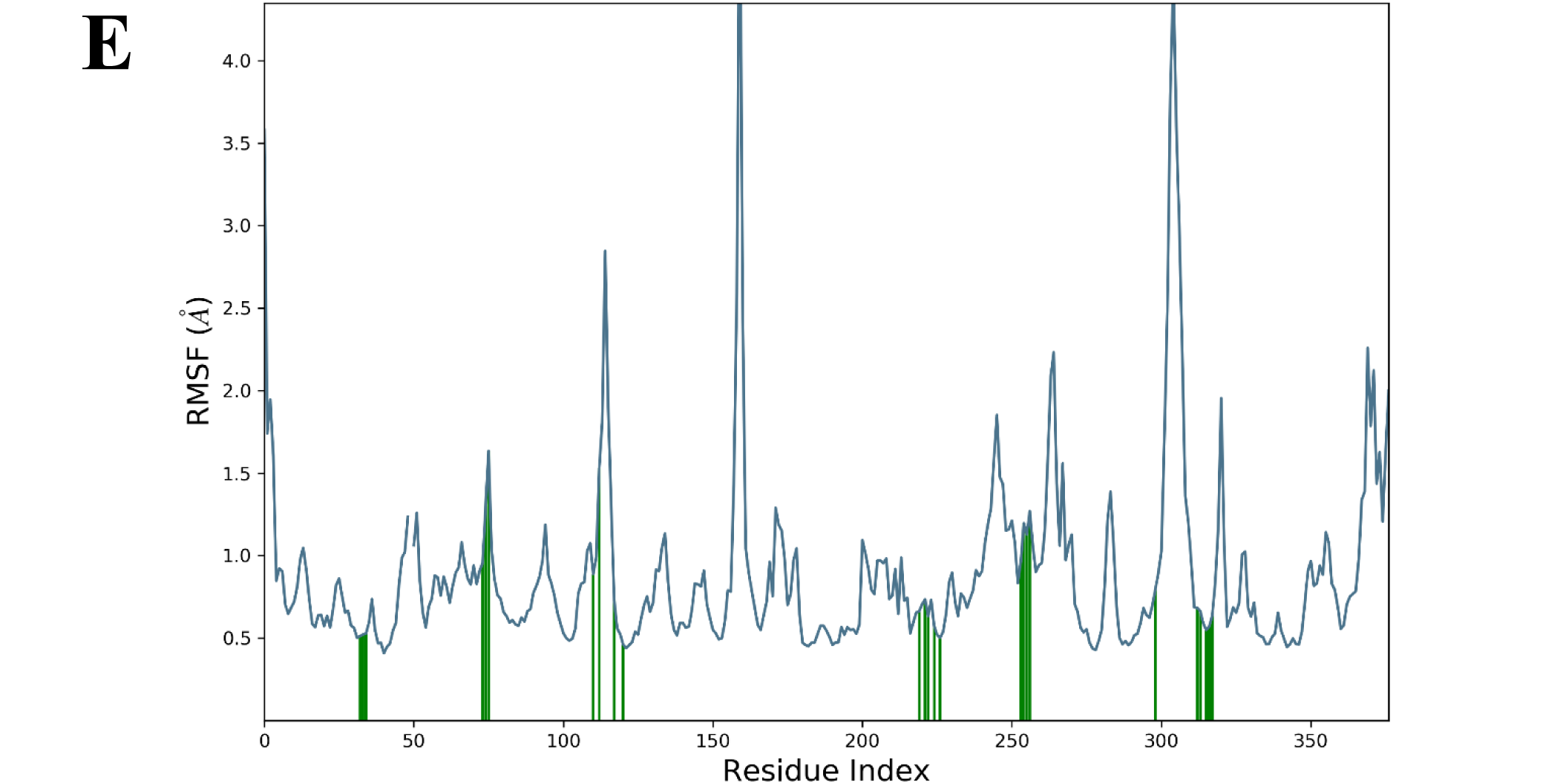
Root Mean Square Fluctuation patterns of β-Secretase bounded to hit compounds. A) Demissidine B) Galantamine C) Solanidine D) Solasidine E) Tomatidine. The green lines show the point of contact of the hit compounds with β-Secretase.

**Figure 7.**
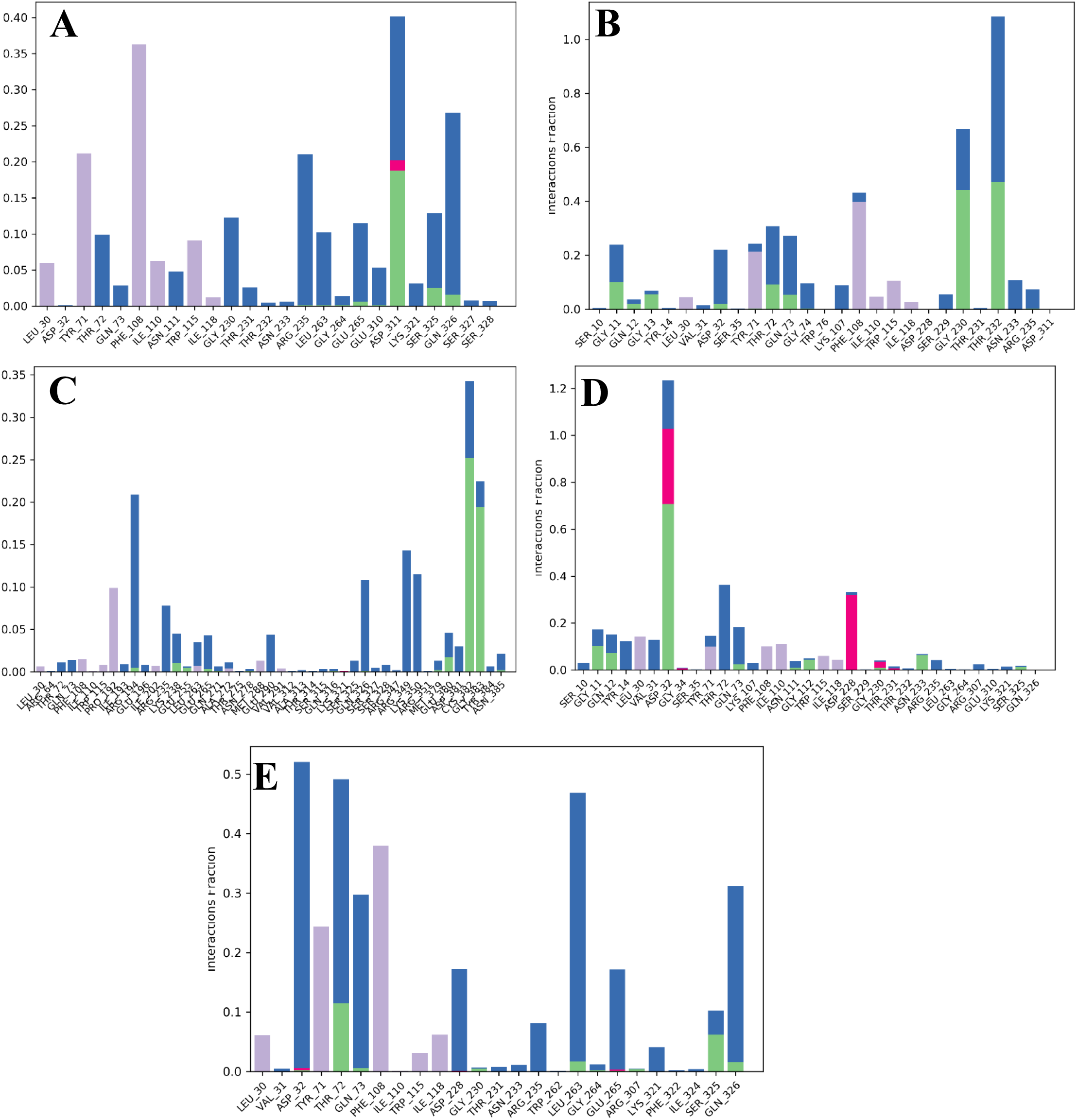
Contact Histogram analysis of β-Secretase bounded to hit compounds. A) Demissidine B) Galantamine C) Solanidine D) Solasidine E) Tomatidine. Green-colored histograms represent hydrogen bonding, purple-colored histograms are for hydrophobic interactions whereas dark blue and red colored histograms are for water bridges and cationic interactions respectively.

**Figure 8.**
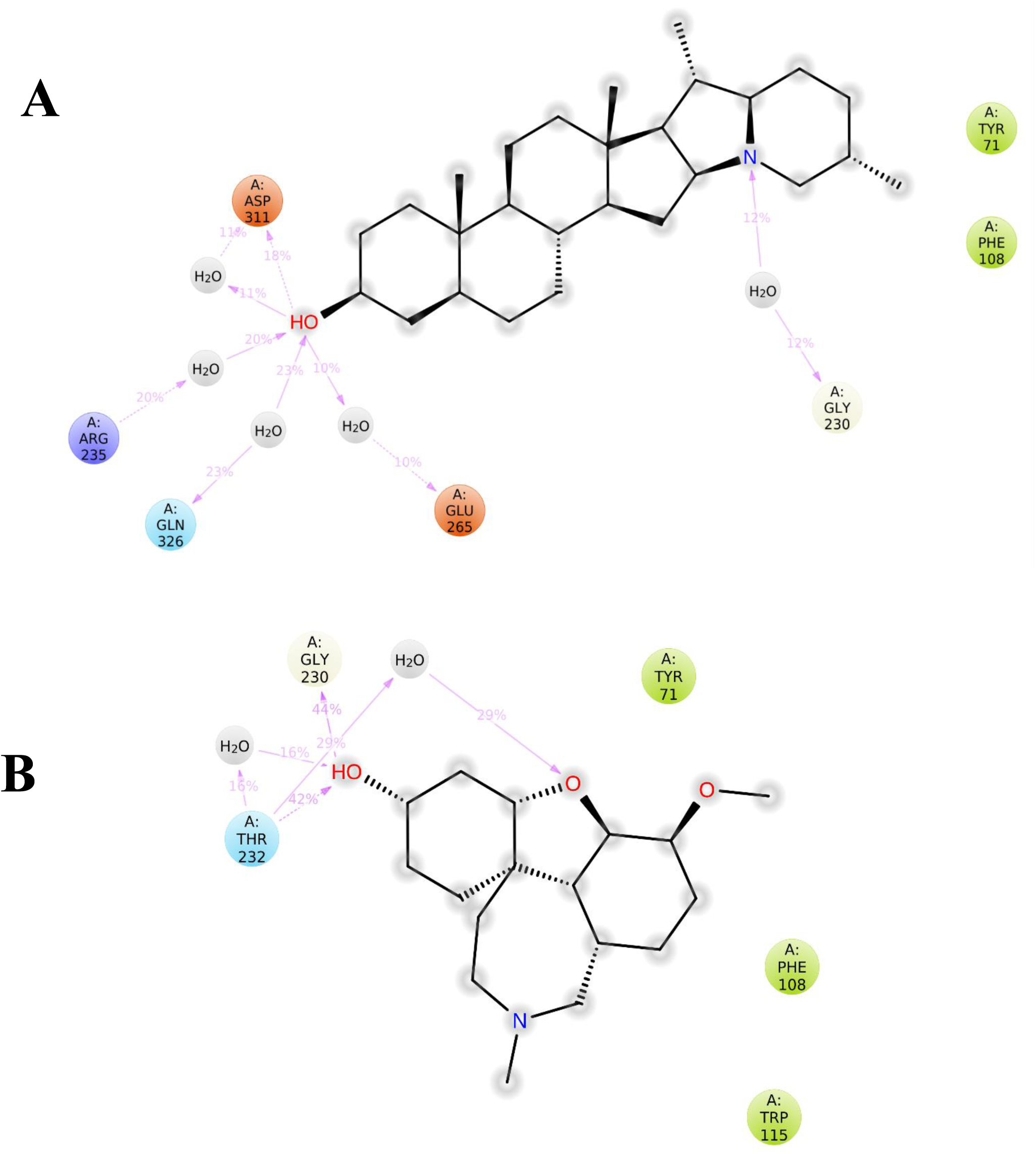
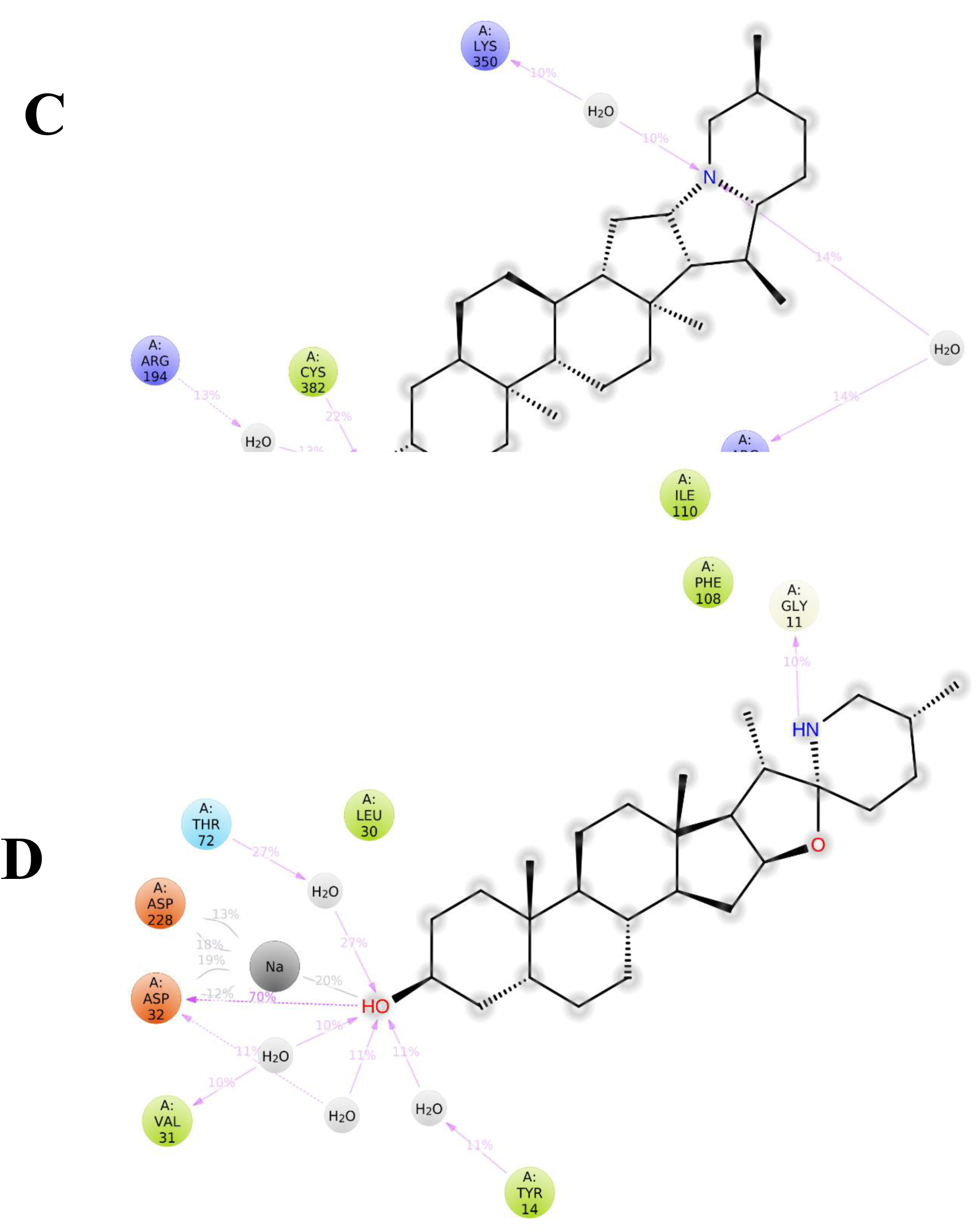
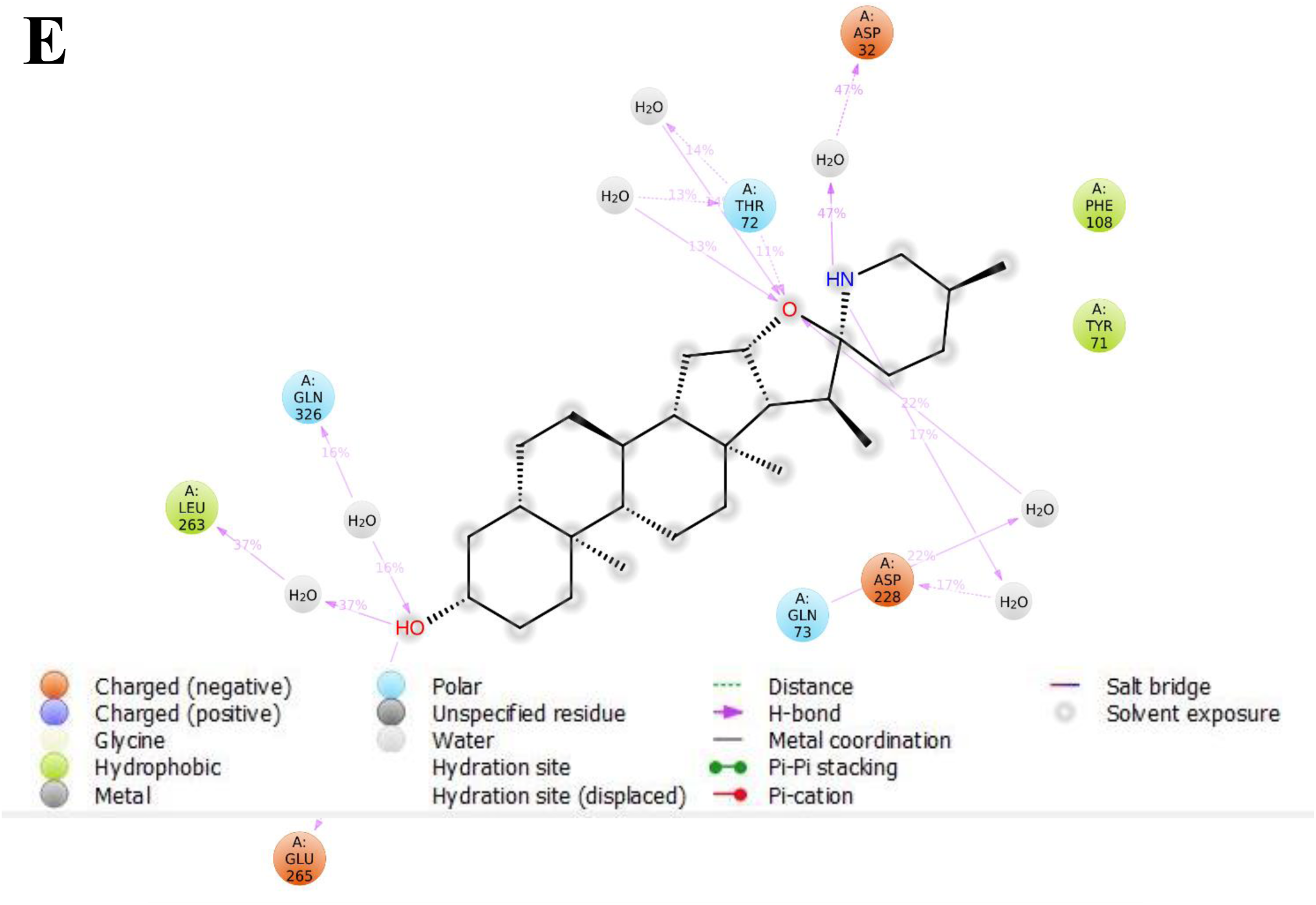
A detailed 2D projection of the atomic interactions that occurred within the selected trajectory (0 through 100 ns). A) Demissidine B) Galantamine C) Solanidine D) Solasidine E) Tomatidine.

**Figure 9.**
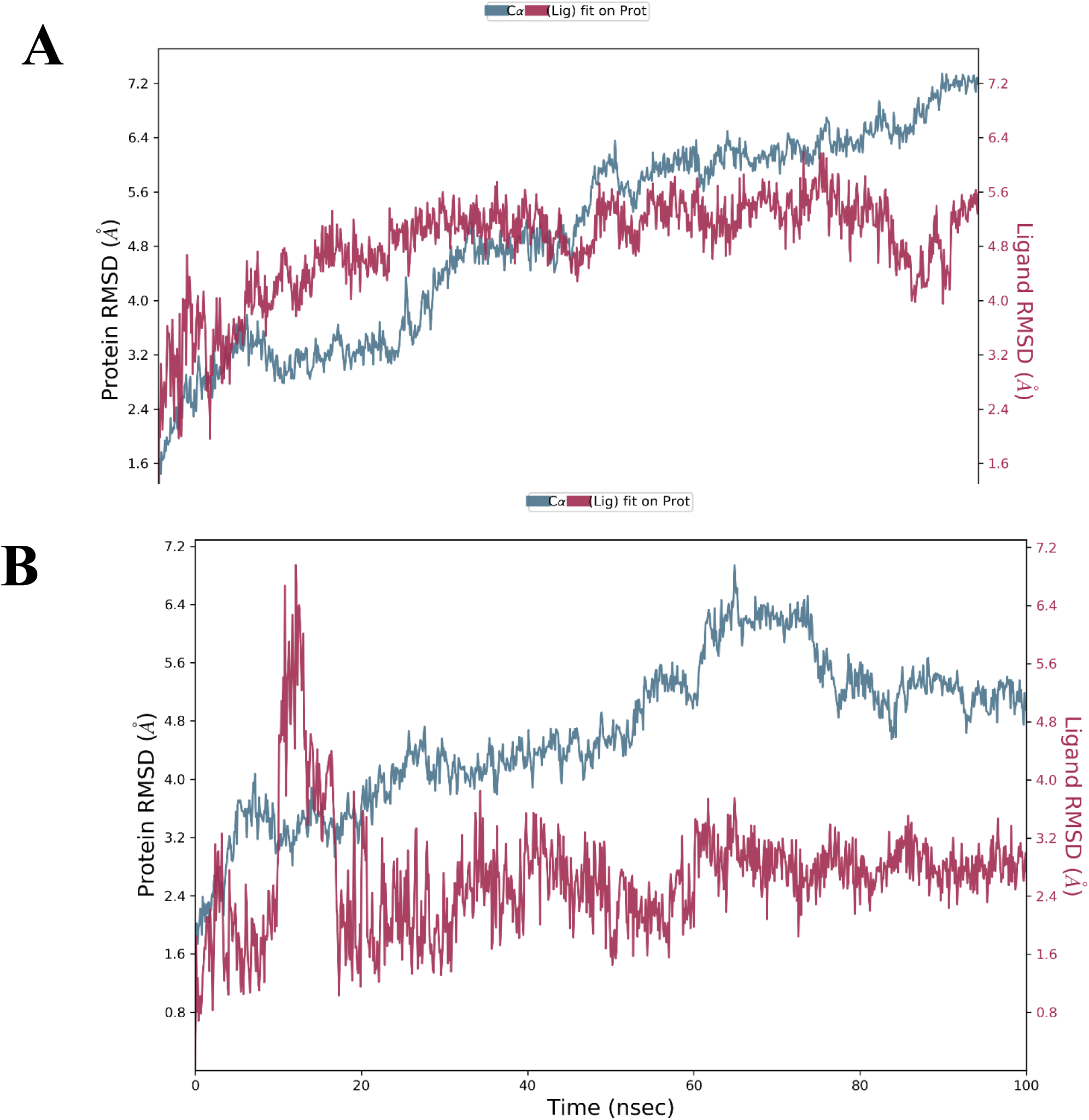
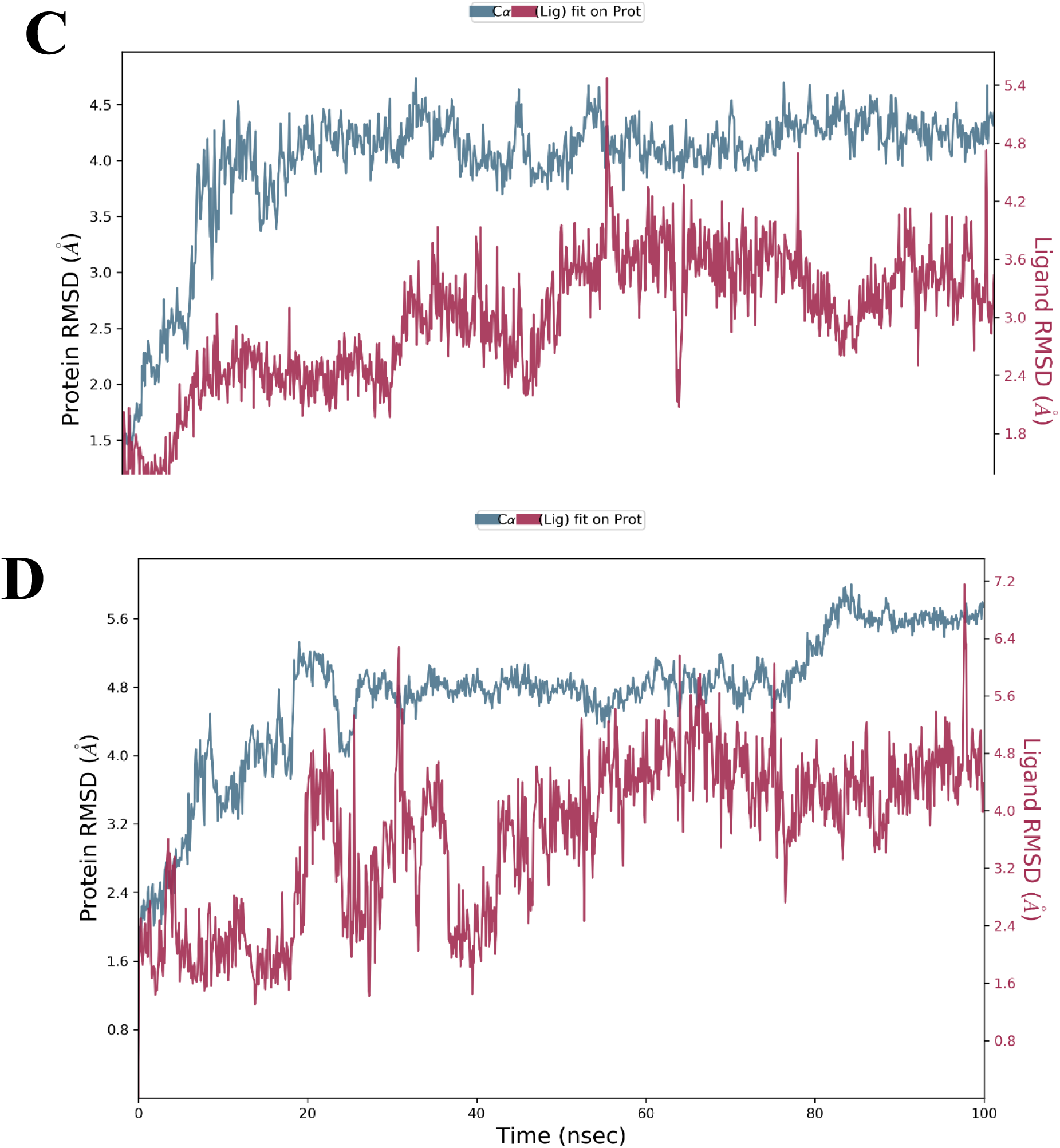
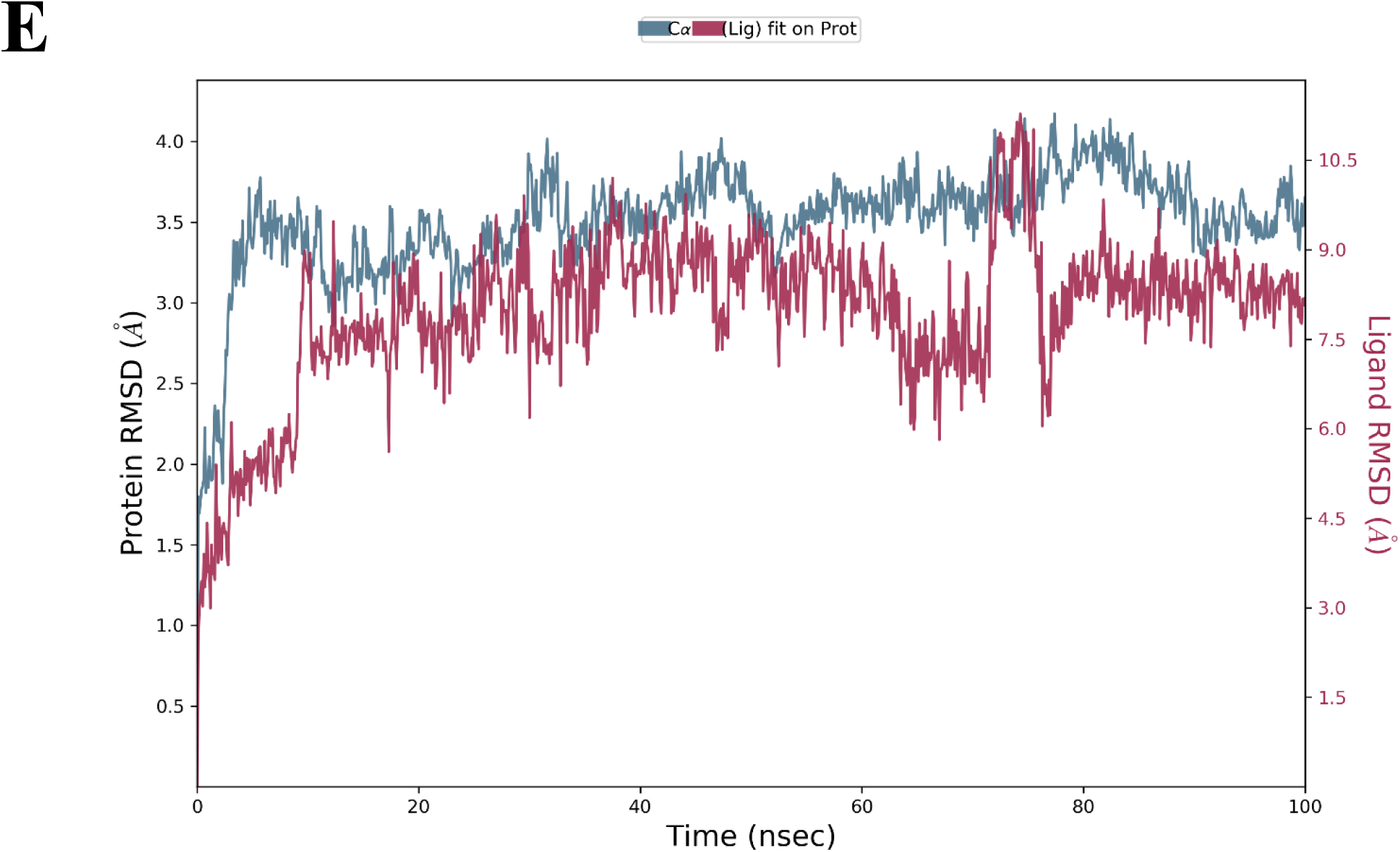
Root Mean Square Deviation of Hit Compound-γ-Secretase complexes. A) Demissidine B) Galantamine C) Solanidine D) Solasidine E) Tomatidine.

**Figure 10.**
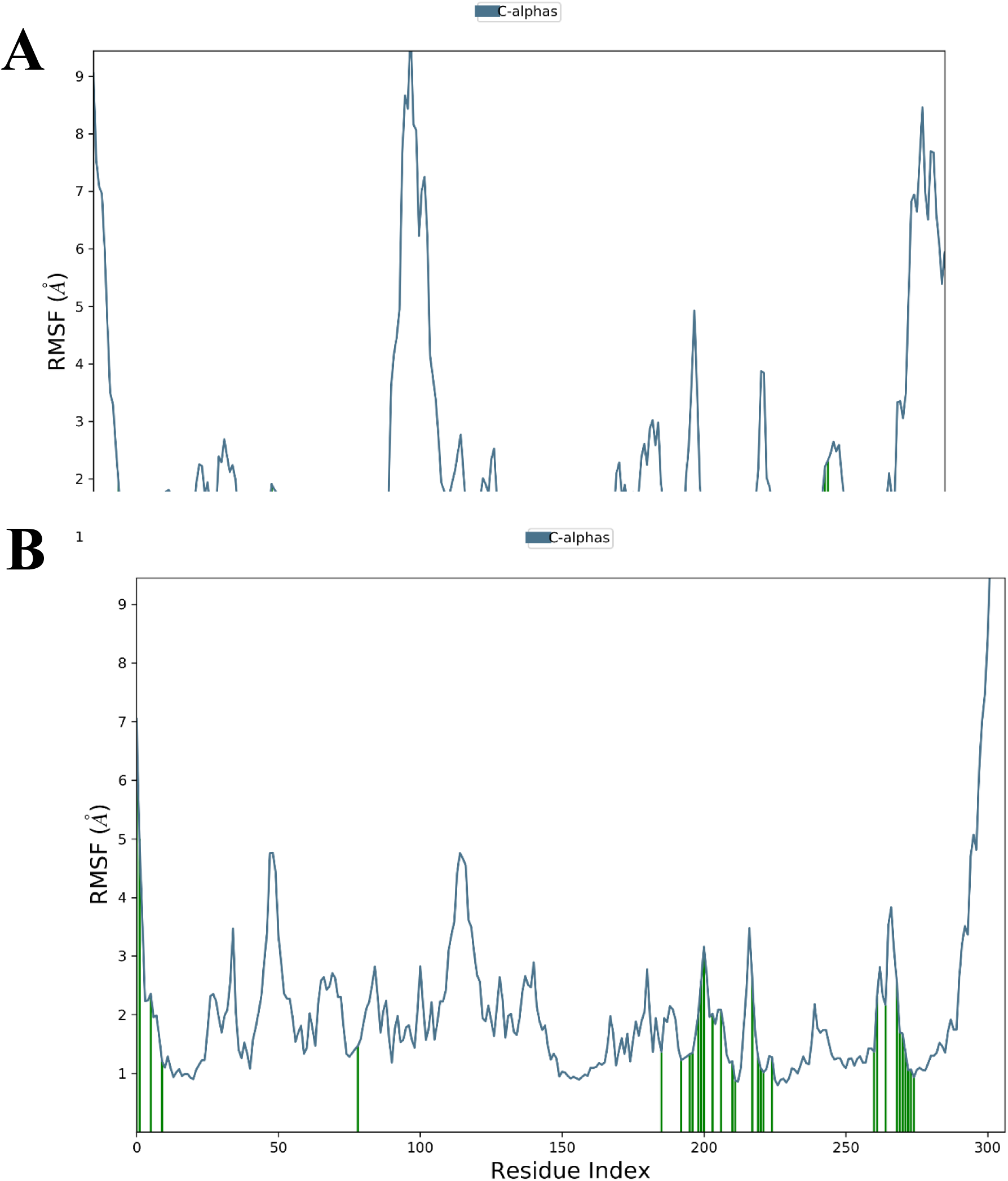
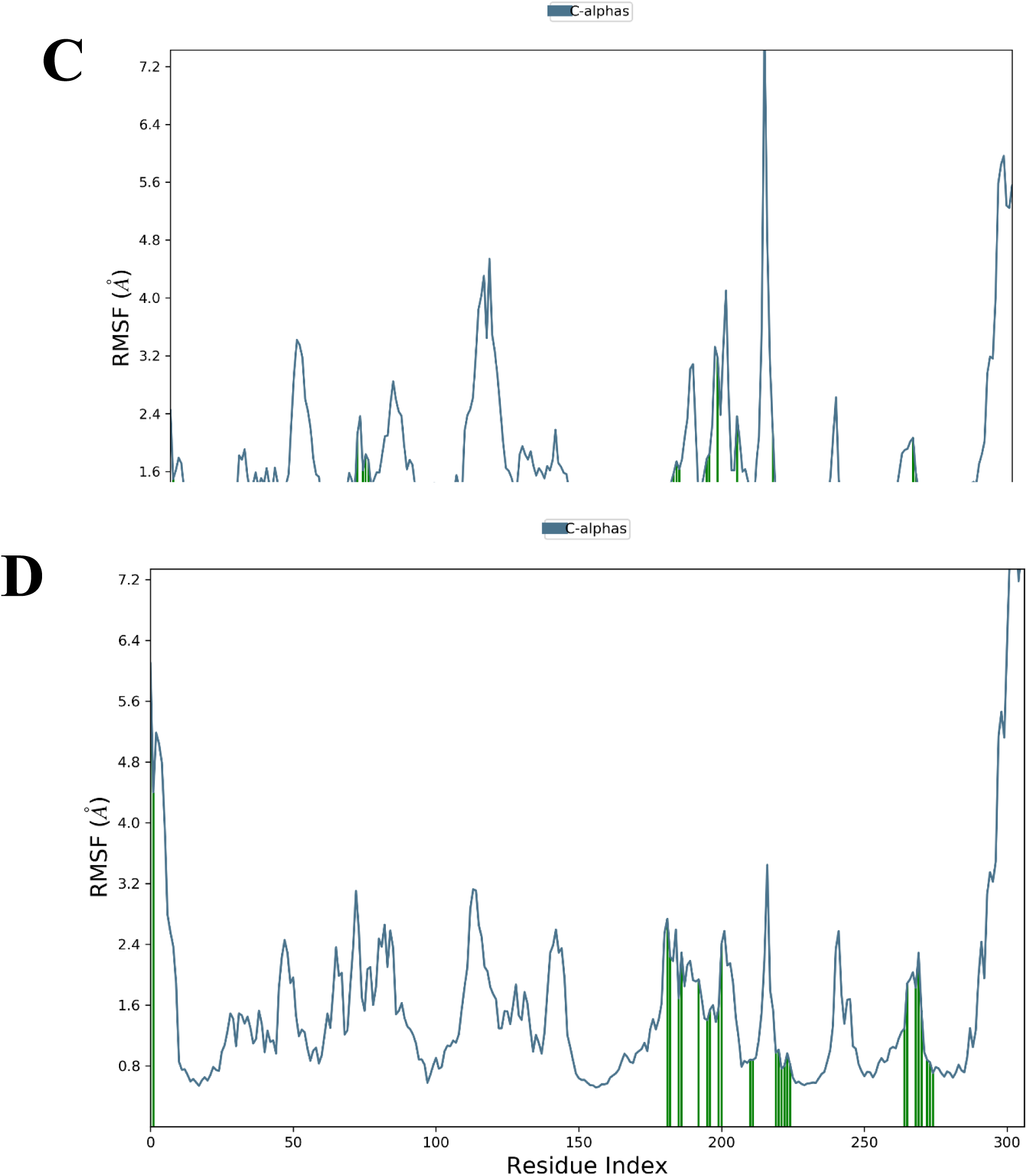
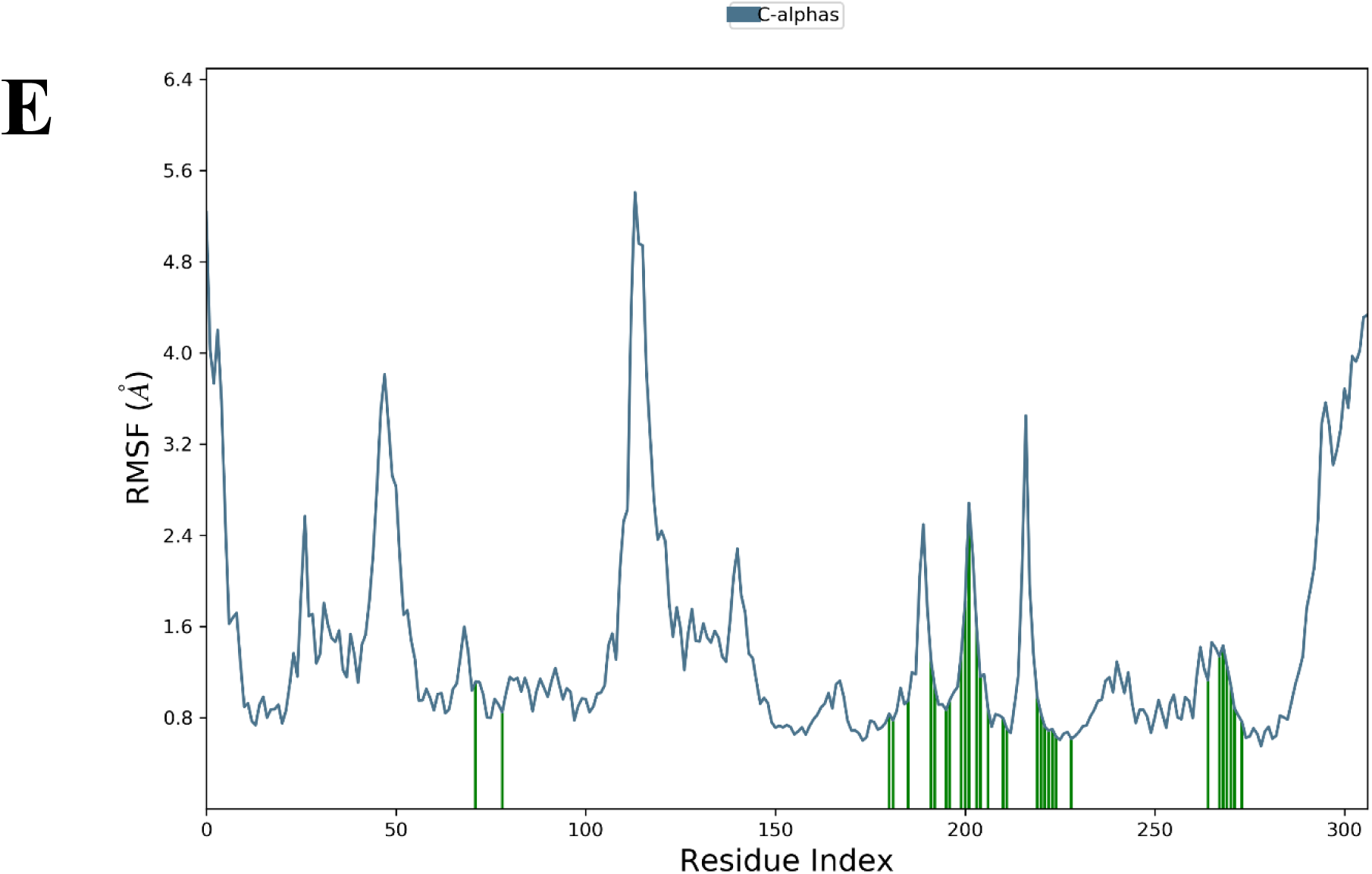
Root Mean Square Fluctuation patterns of γ-Secretase bounded to hit compounds. A) Demissidine B) Galantamine C) Solanidine D) Solasidine E) Tomatidine. The green lines show the point of contact of the hit compounds with β-Secretase.

**Figure 11.**
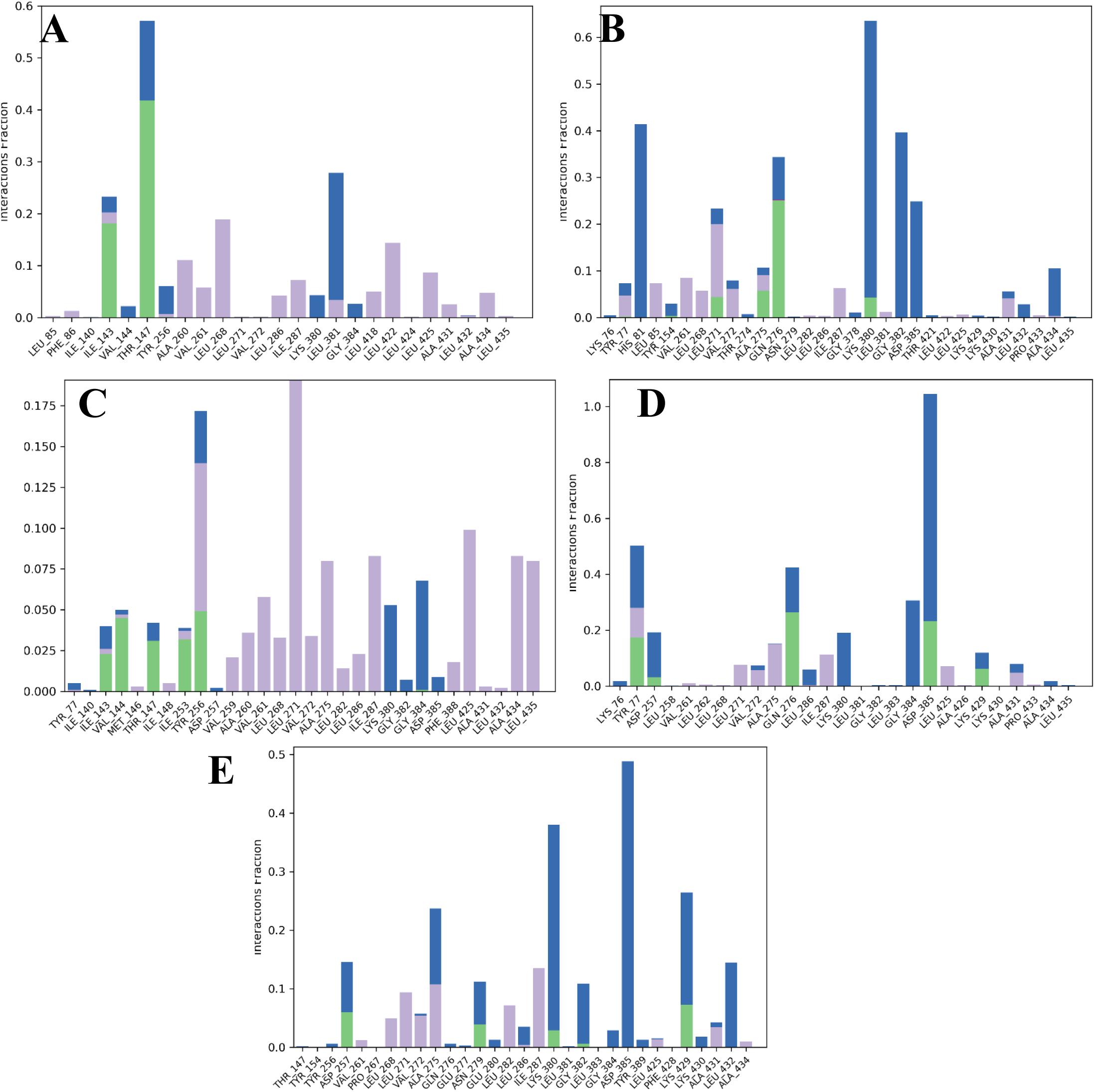
Contact Histogram analysis of γ-Secretase bounded to hit compounds. A) Demissidine B) Galantamine C) Solanidine D) Solasidine E) Tomatidine. Green-colored histograms represent hydrogen bonding, purple-colored histograms are for hydrophobic interactions whereas dark blue and red-colored histograms are for water bridges and cationic interactions respectively.

**Figure 12.**
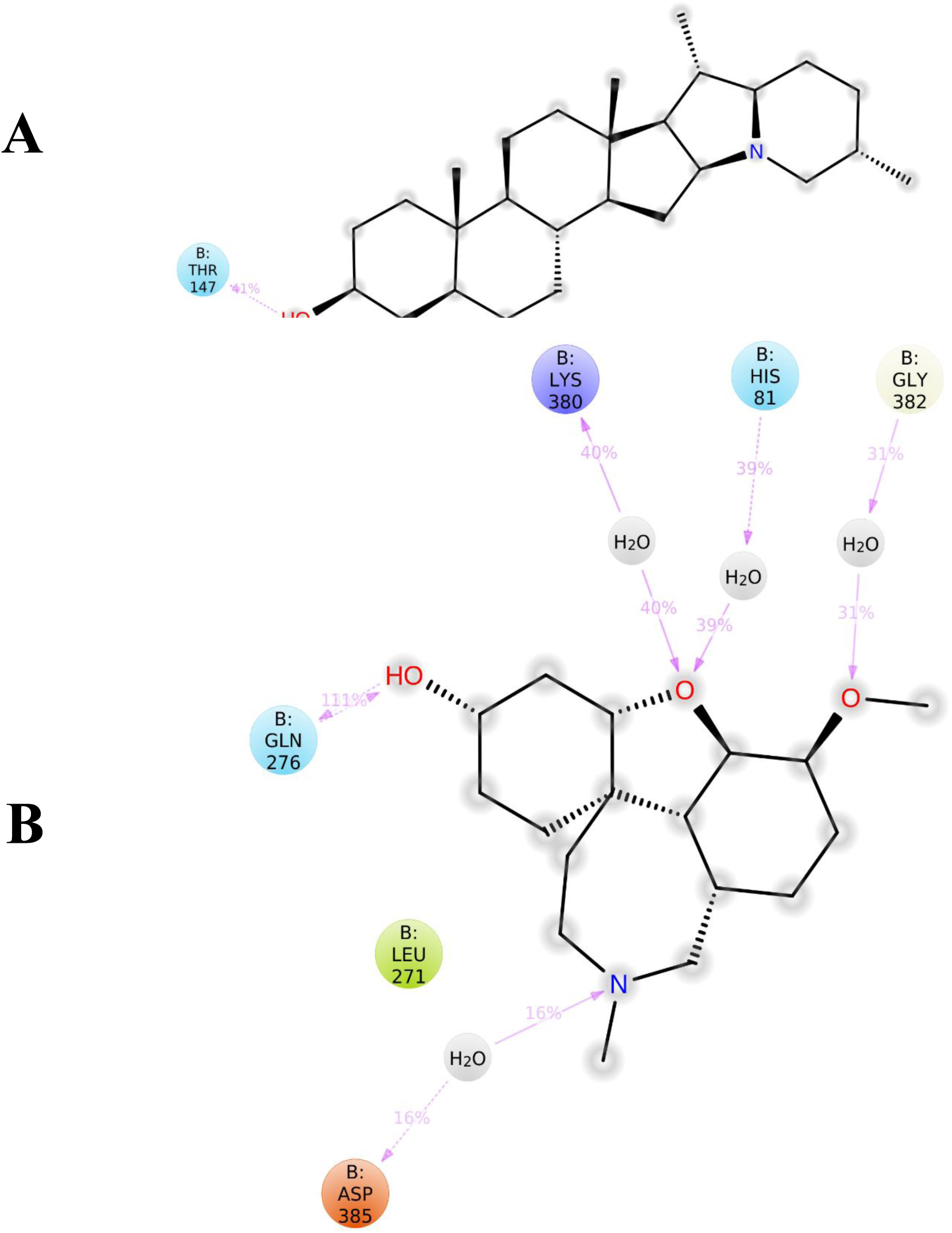
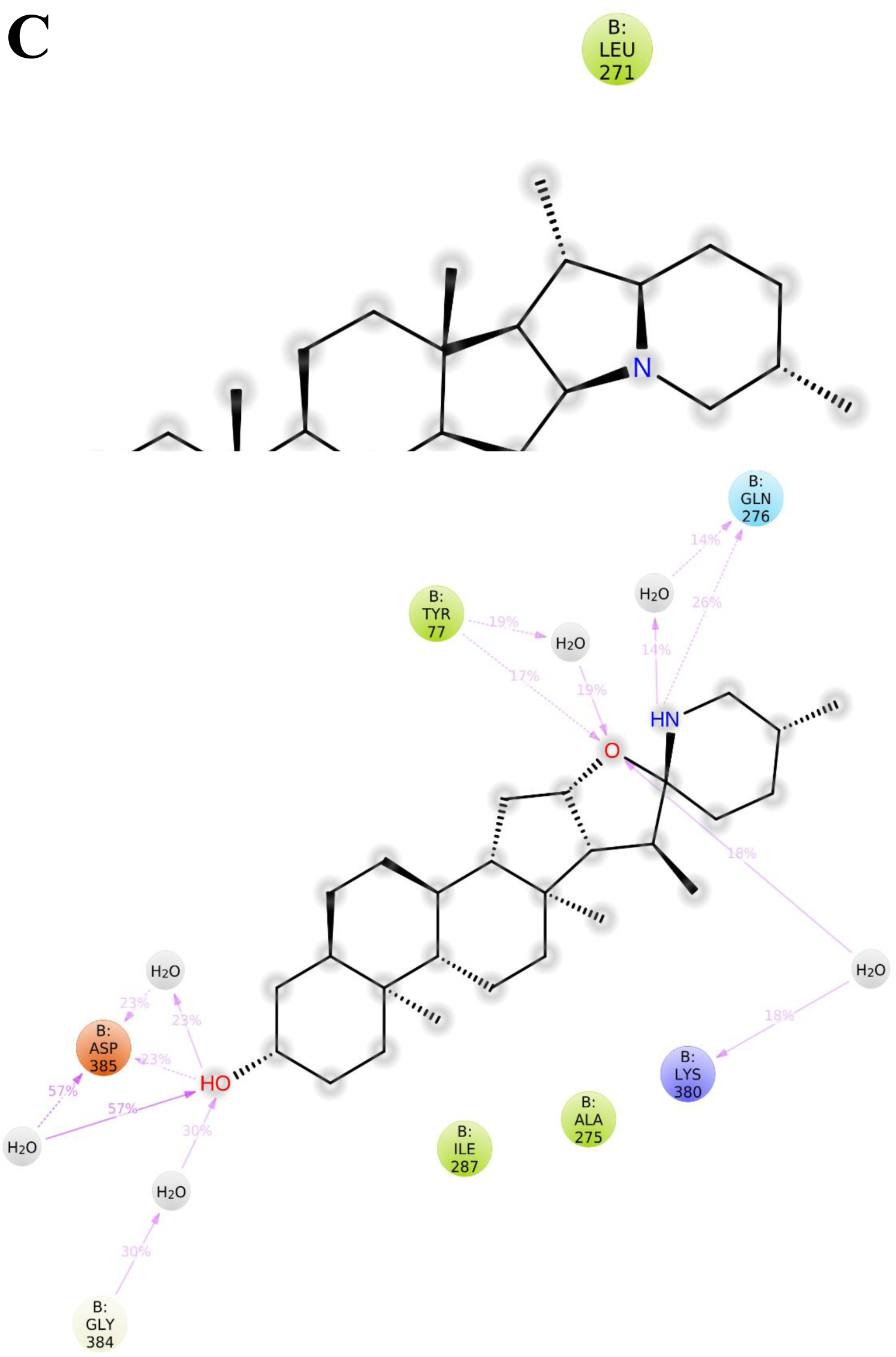
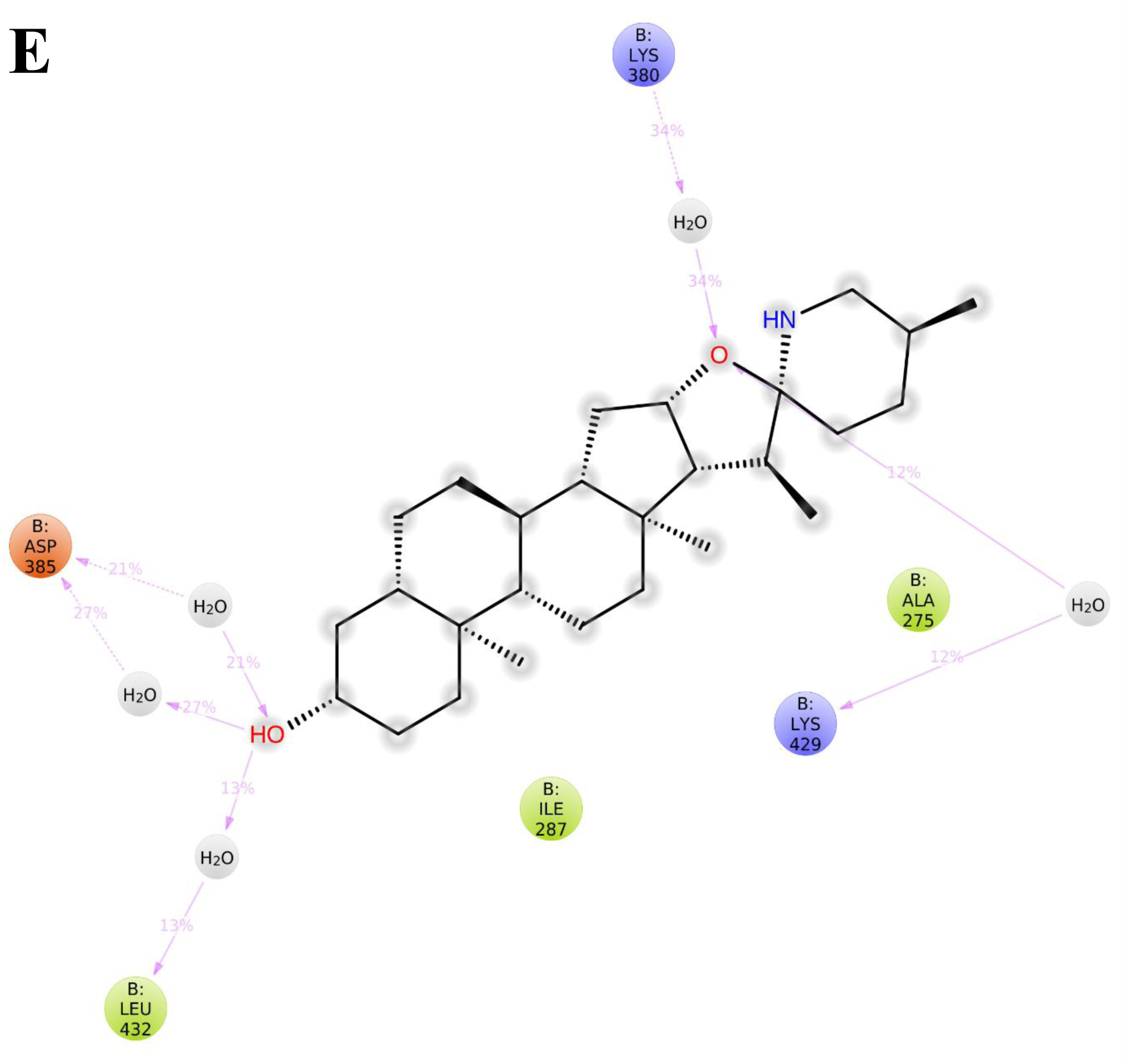
A detailed 2D projection of the atomic interactions that occurred within the selected trajectory (0 through 100 ns). A) Demissidine B) Galantamine C) Solanidine D) Solasidine E) Tomatidine.

**Table 5.** Binding energy (kcal/mol) of hit compounds when docked against secretase using Auto Dock Vina.

| S/No. | Compounds | Binding Energy (kcal/mol) |
| --- | --- | --- |
| 1 | Demissidine | -10.2 |
| 2 | Carpaine | -7.8 |
| 3 | Solasodine | -10.4 |
| 4 | Tomatidine | -11.1 |
| 5 | Solanidine | -10.2 |
| 6 | Galantamine* | -7.2 |
| 7 | L685,458<sup>+</sup> | -9.3 |

\*Control; + co-crystalysed molecule

**Table 6.** *In silico* Pharmacokinetics evaluation through the SwissAdme server.

| Molecule | MW | GI absorp<br>tion | BBB permeant | Pgp substrate | CYP1 A2 inhibitor | CYP2C 19 inhibitor | CYP2 C9 inhibitor | CYP2 D6 inhibitor | CYP3 A4 inhibitor | log Kp (cm/s) |
| --- | --- | --- | --- | --- | --- | --- | --- | --- | --- | --- |
| Carpaine | 478.71 | High | No | No | No | No | No | No | No | -4.75 |
| Demissidine | 399.65 | High | Yes | No | No | No | No | No | No | -3.83 |
| Solanidine | 397.64 | High | Yes | No | No | No | No | No | No | -4.4 |
| Solasodine | 413.64 | High | Yes | Yes | No | No | No | No | No | -5.0 |
| Tomatidine | 415.65 | High | Yes | Yes | No | No | No | No | No | -4.43 |
| Galantamine | 287.35 | High | Yes | Yes | No | No | No | Yes | No | -6.75 |

### 3.2 Discussion

AD has been a great socioeconomic concern that poses an enormous expense to individuals along with their relatives. Several complex cascades of events result from a progressive accumulation of beta-amyloid attributed to factors such as loss of neuronal synapses, neuronal cell death, and progressive neurotransmitter deficits, thus contributing to the clinical symptoms of dementia (Kashyap et al., 2020).

The development of β-secretase and γ-Secretase inhibitors portray an appealing therapeutics potential for Alzheimer’s diseases. β-secretase (BACE1) is a type-1 membrane-anchored aspartyl protease responsible for the processes of β-amyloid peptide in the human brain in patients with Alzheimer’s disease (Maia & Sousa, 2019; Sabbah & Zhong, 2016). The cleavage performed by β-secretase led to the activity of γ-Secretase on the substrate amyloid precursor protein (APP) to generate an intracellular domain (AICD) and a 48- or 49-residue transmembrane peptide (Ab48 or Ab49) (Maia & Sousa, 2019). Accumulation of Aβ peptides of varying lengths and byproducts leads to the formation of β -amyloid plaques, a fingerprint of Alzheimer’s disease (AD). However, the success of some β-secretase and γ-Secretase small molecules inhibitors or modulators has been reported to depict inability to cross the blood-brain barrier, low efficacy or severe side effects (Lei et al., 2015; Ullah et al., 2021). Naturally occurring alkaloids such as Galanthamine and huperzine A have long been used as therapeutic means of providing symptomatic relief to AD patients (Ng et al., 2015). In the current study, the virtual screening strategic step done was to identify selected plant alkaloids that are druggable and could serve as an option in treating AD. Thus, a total of 18 alkaloids exhibited higher affinities when compared to the selected approved drug galantamine and rivastigmine after docking them against β-secretase.

All the selected hit compounds; carpaine, solasodine, tomatidine, solanidin, and demissidine including galantamine from the virtual docking involving β-secretase were subjected to drug-likeness screening. In particular, all the 5 alkaloids show overall good drug-likeness attributes such as low molecular weight (< 500), consensus Log P (CLogP less than 5), number of hydrogen bond acceptors (< 10), and number of hydrogen bond donors (not more than 5) (Table 3). These parameters were set to predict the degree of absorption of the compounds to serve as an oral drug according to the Lipinski Rule of five (R05) (Lipinski et al., 2001). The bioavailability score, pan assay interference compound alert (PAINS alert), leadlikness, synthetic accessibility, and Brenk alerts are considered for the prediction of their medicinal chemistry properties of the 5 selected hit compounds with the control drug (Galantamine) yielded a positive output (Table 4). This predicts the ability of the lead molecules to occur as promiscuous compounds (Baell & Holloway, 2010) related to PAIN’s alert, to deliver druglike potential (Polinsky, 2008), ability to synthesize the hit compounds easily (Ertl & Schuffenhauer, 2009), and structural alert (Brenk et al., 2008).

Thus, this will play a crucial role in the selection process of the compounds for biological evaluation and synthesis of the hit molecules that show a structural-activity relationship with regards to their safety and efficacy during *in vivo* and *in vitro* testing. Noteworthy, the molecular docking conducted with the selected 5 ligands after the first docking period was against the γ-Secretase. This is to have hit compounds that can serve as dual inhibitors of both β-secretase and γ-Secretase. The binding energy scores of the ligand-receptor complex exhibited here were high. Thus, with the dual virtual docking approach in this study, we found out that five ligands; carpaine, solasodine, tomatidine, solanidin, and demissidine are the best inhibitors of β-secretase and γ-Secretase (Table 2 and 5).

It was reported by Bartorelli et al., 2005 that patients with Alzheimer’s disease can benefit from switching to dual inhibitors which help in the stabilization of diseases, reduction in the burden of concomitant psychoactive treatment, and improve the function of cognitive. The major cause of clinical attrition in drug development in the early 1990s was known to be pharmacokinetics and poor drug metabolism (Yang et al., 2019). Early evaluation of absorption, distribution, metabolism, excretion, and toxicity (ADMET) using *in silico* approach helps to predict models for pharmacokinetics of lead compounds to be effective as drugs. Therefore, correct ADMET properties for drug candidates are very important in drug discovery. The *in silico* pharmacokinetics evaluation for all the 5 selected lead molecules was done through the SwissAdme server (Table 6). Gastrointestinal absorption is one of the most important factors to be considered in ADMET properties evaluation because the most frequently used route of drug delivery is the oral delivery system (Ullah et al., 2021) and it is difficult for most oral drugs to cross the intestinal epithelial barrier which determines the bioavailability.

Here, all the 5 selected ligand molecules have a high gastrointestinal absorption rate. Thus, this could increase the efficacy of the drug. Another important ADMET parameter considered is the blood-brain barrier permeability (BBBP) since AD is a neurodegenerative disorder. Therefore, a good drug candidate should be able to cross the blood-brain barrier. All the hit compounds except carpaine were predicted to cross the blood-brain barrier. This show that our lead molecules are promising as the success of some of the reported β-secretase and γ-Secretase inhibitors possess poor blood-brain barrier penetration, low efficacy, and severe side effect, a quick need for wet laboratory experimentation. Williams et al. (2004) reported that the isoforms of CYP enzymes including 1A2, 2C19, 2C9, 2D6, and 3A4 were responsible for about 90% of the oxidative metabolic reactions. Also, the higher a small molecule inhibits these CYP isoforms, the higher the chances of involving in drug-drug interaction with other drugs (Cheng et al., 2011). None of the molecules were predicted to be inhibitors of CYP enzyme isoforms.

Thus, this shows that these drug candidates can be used alone or with other drugs without resulting in any adverse effects or therapeutic failure. P-glycoproteins are proteins involved in transporting many drugs through the cell membrane and they are very important in intestinal absorption, brain penetration, drug metabolism, and excretion (Broccatelli et al., 2011; Ullah et al., 2021). In addition, P-gp substrate or inhibition may affect the bioavailability and safety of drugs inside the cell. In this study, we included Pgp substrate in here as it helps to minimize the cost and time with integration with *in vitro* evaluation. However, solasodine and tomatidine were predicted to none-substrate. This shows that these two compounds may have the potential of overcoming multidrug resistance (Broccatelli et al., 2011).

Our study showed that only solasodine bound with β-secretase by forming interaction with one of the key amino acids among the catalytic dyad residue, Asp 32 (Fig G ii). This indicates an important contribution to the cleavage of the amyloid precursor protein. It is also noteworthy, that the Tyr 71 residue contributes to the inhibitor binding at the active site by changing its conformation (Barman & Prabhakar, 2014; Sabbah & Zhong, 2016). Carpaine was able to exhibit 8 interactions with the amino acid residue within the binding cavity of β-secretase; Thr72, Tyr71, Gly230, Leu30, Trp115, Phe108, Gln73, and Ile110. Also, it formed 12 hydrophobic interactions within the binding portion of γ-Secretase; Leu166, Leu286, Gly382, Asp257, Leu 268, Ile143, Ile253, Gly384, Phe388, Ile387, Leu383, and Trp165. Demissidine contributes one hydrogen bond with both β-secretase and γ- Secretase; Ser325 and Gln276 respectively. Similarly, demissidine established 7 and 9 hydrophobic interactions with β-secretase and γ-Secretase: Ile110, Phe108, Leu30, Trp115, Tyr71, Gln73, Thr72; and Gly382, Pro433, Val261, Ala431, Lys380, Leu282, Val272, Ile287, and Asp385 respectively.

Phe108, Trp115, Leu30, Gln12, Ile110 interact with galantamine hydrophobically, and Gln73 via hydrogen bond with β-secretase; while Leu381, Ile287, Val272, Lys380, Leu425, and Ala430 interact with galantamine through Hydrophobic interaction; and Gly382 and Leu432 with hydrogen bond interaction for γ-Secretase. Solanidine established hydrophobic interaction with Phe108, Tyr71, Leu30, Trp115, Gln73, and 1 hydrogen bond interaction with Gln276 amino acid residues of β-secretase; while Val261, Pro433, Ala431, Lys380, Leu282, Val272, Ile287, Gly382, and Asp385 formed hydrophobic interaction in the catalytic domain of γ-Secretase, in addition to one hydrogen bond with Gln276. Hydrophobic bond interaction was observed with Four (4) and nine (9) amino acid residues within the catalytic domain of β-secretase and γ-Secretase for solasodine; Tyr71, Ile118, Asp32, and Ile110; and Asp385, Gly382, Lys380, Ala431, Val272, Leu282, Ile287, Leu268, Gly384 respectively. However, hydrogen bond interaction with solasodine was established only with β-secretase through Gly11.

Tomatidine formed hydrophobic bond interaction with the amino acid residue of β-secretase; Gly230, Thr 231, Thr 232, Asn 233, Tyr71, and Thr72, while hydrophobic bond interaction was established with Gly384, Pro433, Ala431, Lys380, Leu282, Val272, Ile287, Leu268, and Gly382; and hydrogen bond interaction with Asp385 was formed with the amino acid residue within the catalytic domain of γ-Secretase. It is imperative to know that Solasodine, Tomatidine, Solanidin, and Demissidine are *Solanum* alkaloids. *Solanum* alkaloids have been a topic of interest since decades. They were reported to have exhibited promising biological potentialities including anticancer activities, neurotoxicity and neuroprotective activity, anti-inflammatory effects, and antimicrobial activity both *in vitro* and *in vivo*. However, these best molecules were found to interact with key amino acids in the active site of β and γ secretase and were predicted to be safe. Therefore, these lead molecules can be investigated further to develop potent and safe anti-AD drugs.

## Conclusion

Finally, phytocompounds, especially alkaloids, were known for their therapeutic value applications, sometimes considered to reduce Alzheimer’s disease (Ng et al., 2015; Rao et al., 2012). In this current study, we analyzed 54 alkaloids in stepwise computational virtual docking approaches, pharmacokinetics evaluation, drug-likeness prediction, and medicinal chemistry evaluation to identify potential dual inhibitors of β-secretase and γ-Secretase with good efficacy and safety. Ultimately, 5 ligands- Carpaine, Solasodine, Tomatidine, Solanidin, and Demissidine were selected as the best inhibitors. However, there will be an urgent need for *in vitro* and *in vivo* studies with the appropriate model to give more insight into the hit compounds.

## Notes

### Competing Interest Statement

The authors have declared no competing interest.

